# Molecular basis of human nuclear and mitochondrial tRNA 3’-processing

**DOI:** 10.1101/2024.04.04.588063

**Authors:** Arjun Bhatta, Bernhard Kuhle, Ryan D. Yu, Lucas Spanaus, Katja Ditter, Katherine E. Bohnsack, Hauke S. Hillen

**Affiliations:** University Medical Center Göttingen, Department of Cellular Biochemistry, Humboldtallee 23, 37073 Göttingen, Germany; Max Planck Institute for Multidisciplinary Sciences, Research Group Structure and Function of Molecular Machines, Am Fassberg 11, 37077 Göttingen, Germany; University Medical Center Göttingen, Department of Molecular Biology, Humboldtalle 23, 37073 Göttingen, Germany; Cluster of Excellence “Multiscale Bioimaging: from Molecular Machines to Networks of Excitable Cells” (MBExC), University of Göttingen, D-37075 Göttingen, Germany; Göttingen Center for Molecular Biosciences (GZMB), Research Group Structure and Function of Molecular Machines, University of Göttingen, D-37077 Göttingen, Germany

**Author notes:** These authors contributed equally (Arjun Bhatta, Bernhard Kuhle, Ryan D. Yu).

## Abstract

Eukaryotic transfer RNA (tRNA) precursors undergo sequential processing steps to become mature tRNAs. In humans, ELAC2 carries out 3’-end processing of both nucleus-encoded (nu-tRNAs) and mitochondria-encoded tRNAs (mt-tRNAs). ELAC2 is self-sufficient for processing of nu-tRNAs, but requires TRMT10C and SDR5C1 to process most mt-tRNAs. Here, we show that TRMT10C-SDR5C1 specifically facilitate processing of structurally degenerate mt-tRNAs lacking the canonical elbow. Structures of ELAC2 in complex with TRMT10C, SDR5C1 and two divergent mt-tRNA substrates reveal two distinct mechanisms of pre-tRNA recognition. While canonical nu-tRNAs and mt-tRNAs are recognized by direct ELAC2-RNA interactions, processing of non-canonical mt-tRNAs depends on protein-protein interactions between ELAC2 and TRMT10C. These results provide the molecular basis for tRNA 3’-processing in both the nucleus and mitochondria and explain the organelle-specific requirement for additional factors. Moreover, they suggest that TRMT10C–SDR5C1 evolved as a mitochondrial tRNA maturation platform to compensate for the structural erosion of mt-tRNAs in bilaterian animals.

## INTRODUCTION

Transfer RNAs (tRNAs) are key mediators between the RNA and protein worlds that decode genetic information in mRNAs to sequences of amino acids during translation^1^. They are transcribed as precursor transcripts (pre-tRNAs), which need to be post-transcriptionally processed and matured to become functional^2–5^. The first steps of tRNA processing involve removal of leader- and trailer sequences at their 5’- and 3’-ends^5–8^. These universal steps are followed by species- and organelle-specific maturation steps, such as 3’-CCA-addition, chemical modifications, and removal of introns by tRNA-splicing^2,4,5,7,9–12^. These steps are crucial for the proper function of tRNAs, and mutations in tRNA-maturation factors can cause severe diseases in humans ^11,13–15^.

Consistent with their ancient evolutionary origin and fundamental role in protein biosynthesis, the basic “canonical„ structure of tRNAs is highly conserved^16–18^. They adopt a typical cloverleaf secondary structure comprised of four arms (anticodon, acceptor, T- and D arm), which fold into an L-shaped tertiary structure^19–21^. Its opposing ends are formed by i) the acceptor arm, which contains the 5’- and 3’-ends of the tRNA and serves as the site of aminoacylation, and ii) the anticodon loop, which contains the eponymous anticodon-triplet for decoding mRNA codons during ribosomal translation^20,21^. The structural core of canonical tRNAs consists of a tight network of tertiary interactions between the D- and T-loops, forming the characteristic “tRNA elbow„ and stabilizing the overall L-shaped fold^21,22^. These conserved structural features serve as recognition sites for enzymes that carry out tRNA maturation and charging^23,24^.

In addition to about 300 different cytoplasmic tRNAs encoded by ∼500 tRNA genes in the human nuclear genome^25,26^, the human mitochondrial genome (mtDNA) encodes a minimal set of 22 tRNAs (mt-tRNAs), which are required for the synthesis of the 13 mtDNA-encoded essential subunits of the oxidative phosphorylation complexes^27^. Despite their key role in cellular energy homeostasis, animal mt-tRNAs underwent a unique process of sequence and structural erosion, leading to the loss of many of the conserved structural features required for nu-tRNA processing, modification, and aminoacylation^28–30^. Consequently, the enzymes performing these steps are either specialized in recognizing only nuclear or only mitochondrial tRNAs or evolved unique mechanisms to serve tRNAs from both genomes^31–35^.

The first steps of tRNA processing are carried out by the endoribonucleases (RNases) P and Z, which cleave the 5’-leader and 3’-trailer sequences from primary pre-tRNA transcripts^4,10,36^. In humans, nu-tRNAs and mt-tRNAs are processed at the 5’-end by two different RNase Ps of distinct architectures and evolutionary origins^32,37,38^. Nu-tRNAs are processed by a large ribonucleoprotein complex (nu-RNase P), which recognizes the acceptor–T-arm domain and the elbow of its pre-tRNA substrate^39^. By contrast, mt-tRNAs are processed by a protein-only RNase P composed of the methyltransferase TRMT10C (MRPP1), the dehydrogenase SDR5C1 (MRPP2/HSD17B10) and the endoribonuclease PRORP (MRPP3)^32^, which recognizes pre-tRNA substrates via their overall L-shaped structure and conserved elements in the anticodon loop^40^. These interactions are mediated by the TRMT10C–SDR5C1 subcomplex, which has been proposed to act as a maturation platform for multiple steps of mt-tRNA processing^40,41^.

In contrast to 5’-end processing, 3’-end processing of both human nu-tRNAs and mt-tRNAs is catalyzed by a common RNase Z enzyme, named ELAC2^31,42^. RNase Z enzymes belong to the β-lactamase family of metal-dependent endonucleases^43–45^. The β-lactamase domain contains the active site, with two catalytic zinc ions (Zn^2+^) coordinated by a conserved HXHXDH motif, and an RNase Z-specific sequence insertion called the “flexible arm„, which binds the elbow of the pre-tRNA substrate^44–47^. Two types of RNase Z enzymes exist: a ubiquitous short form (RNase Z_S_) that contains a single β-lactamase domain that assembles into homodimers, and a eukaryote-specific long form (RNase Z_L_) with two beta-lactamase domains that presumably evolved by gene duplication and fusion^48^. ELAC2 is an RNase Z_L_that localizes to both the nucleus and mitochondria^31,49^. In the nucleus, ELAC2 by itself is sufficient for tRNA processing, as it can efficiently cleave nu-tRNA precursors without additional factors^50^. In contrast, *in vitro* studies show that ELAC2 requires TRMT10C and SDR5C1 for efficient processing of most mt-tRNAs, suggesting that human mitochondrial RNase Z may be a multi-subunit complex^41^. To date, no structures of ELAC2 or other RNase Z_L_enzymes in complex with their substrate are available, and the structural and mechanistic basis of eukaryotic tRNA 3’-processing remains elusive. Furthermore, it is not known how ELAC2 specifically recognizes its structurally diverse nuclear and mitochondrial tRNA substrates, why the TRMT10C–SDR5C1 subcomplex is required for processing of most mt-tRNAs, and how the strict order of tRNA 5’-processing followed by 3’-processing is ensured.

Here, we use a combination of *in vitro* biochemical assays and single-particle cryogenic electron microscopy (cryo-EM) to elucidate the mechanism of human tRNA 3’-processing. Using a reconstituted system, we show that TRMT10C–SDR5C1 specifically facilitates processing of structurally degenerate mt-tRNAs with reduced elbow regions, suggesting that it compensates for the absence of this otherwise-conserved element. Cryo-EM structures of ELAC2 in complex with TRMT10C–SDR5C1 and either a canonical (mt-tRNA^Gln^) or degenerate mt-tRNA precursor (mt-tRNA^Tyr^) reveal how ELAC2 binds and accurately processes these structurally divergent tRNAs, providing a mechanistic model for tRNA 3’-end processing in both the nucleus and mitochondria. They show that TRMT10C stabilizes the tertiary fold of degenerate mt-tRNAs and facilitates ELAC2 binding via direct protein-protein interactions, thereby compensating for the loss of protein-RNA interactions with the conserved elbow structure in degenerate mt-tRNAs. Finally, our data suggest a rationale for the strict sequence of tRNA processing steps. Taken together, these results provide a molecular picture of human tRNA 3’-processing in the nucleus and mitochondria, explain the requirement of TRMT10C and SDR5C1 for mitochondrial tRNA processing, and yield insights into the evolution of the TRMT10C–SDR5C1 complex as the mitochondrial tRNA maturation platform.

## RESULTS

### The TRMT10C–SDR5C1 complex facilitates 3’-end processing of structurally degenerate mt-tRNAs

To investigate the mechanism of human tRNA 3’-processing, we first set out to understand how a single ELAC2 enzyme can process structurally highly diverse nuclear and mitochondrial pre-tRNA substrates. We started by re-examining previous data by Reinhard et al. on the differential dependence of ELAC2 on TRMT10C– SDR5C1 for *in vitro* processing of mt-tRNAs^41^ (**Figure 1A, B**). In particular, we compared the degree to which ELAC2-mediated cleavage was reported to be dependent on TRMT10C–SDR5C1 (strong, intermediate and no dependence) to the structural properties of the respective mt-tRNAs^28,41,51,52^. This revealed that the four mt-tRNAs exhibiting “no dependence„ on TRMT10C– SDR5C1 are the only mt-tRNAs predicted to form nu-tRNA-like canonical or near-canonical tertiary elbow interactions. By contrast, mt-tRNAs with highly reduced D- and T-loops and predicted to lack tertiary elbow interactions consistently showed “strong dependence„ on TRMT10C–SDR5C1. This suggests that the differential dependence of tRNAs on TRMT10C– SDR5C1 for 3’-processing by ELAC2 may be related to the structural properties of the tRNA elbow region.

**Figure 1.**
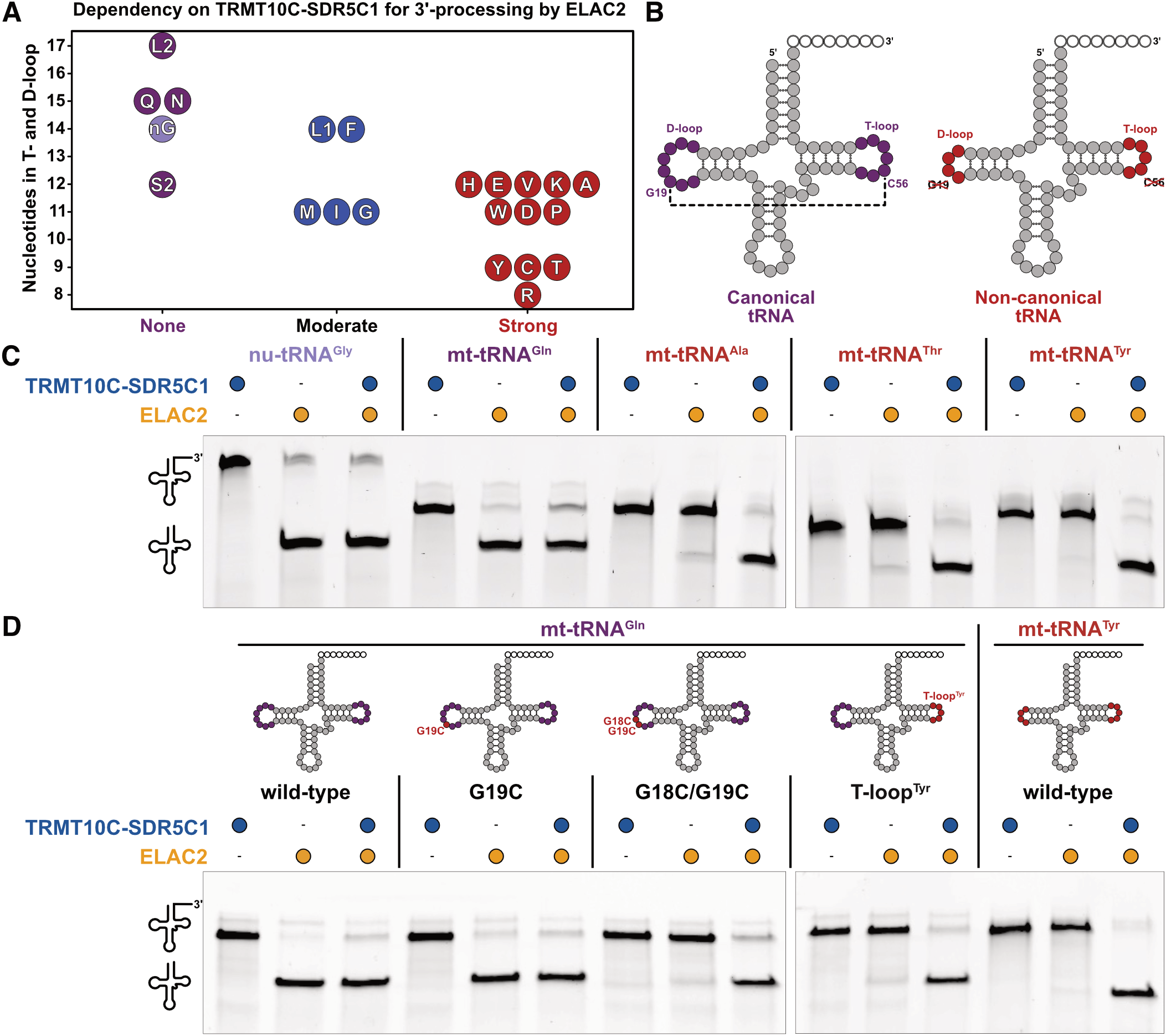
Differential dependence of nuclear and mitochondrial tRNAs on TRMT10C–SDR5C1 for 3’-processing by ELAC2. **A)** The dependency of mt-tRNAs on TRMT10C–SDR5C1 for 3’-processing by ELAC2 correlates with the degeneracy of their elbow region. Mt-tRNAs are labelled with the one-letter code for their cognate amino acid. Mt-tRNA^Leu^(UUR), mt-tRNA^Leu^(CUN) and mt-tRNA^Ser^(UCN) are labelled as L1, L2 and S2, respectively. Mt-tRNA^Ser^(AGY) is not included, as it is not processed efficiently by ELAC2 either in presence or absence of TRMT10C–SDR5C1^41,71^. The sum of nucleotides in the D- and T-loops of each mt-tRNA is plotted against their dependency on TRMT10C–SDR5C1 according to Reinhard et al^41^. For reference, the human nuclear tRNA^Gly^ is included (nG), which shows no dependence on TRMT10C–SDR5C1 **(Figure 1C)**. **B)** Secondary structures of mt-tRNA^Gln^ (left) and mt-tRNA^Tyr^ (right), representing a canonical and degenerated cloverleaf fold, respectively. While mt-tRNA^Gln^ can form canonical D-loop-T-loop tertiary interactions including the conserved G19-C56 WC-pair (indicated by dashed lines), mt-tRNA^Tyr^ lacks conserved nucleotides to form tertiary interactions. **C)** *In vitro* cleavage assays showing the differential dependency between canonical and degenerated tRNAs on TRMT10C– SDR5C1 for 3’-processing by ELAC2. **D)** *In vitro* cleavage assays demonstrating the gain of TRMT10C–SDR5C1-dependence by mt-tRNA^Gln^ upon introduction of mutations disrupting the canonical elbow structure. Gels in **C** and **D** are representatives of three independent replicates.

To test this hypothesis, we reconstituted a 3’-processing system using purified recombinant ELAC2, TRMT10C and SDR5C1 as well as *in vitro* transcribed pre-tRNA substrates containing either canonical (nu-tRNA^Gly^ and mt-tRNA^Gln^) or highly degenerated tRNA elbow structures (mt-tRNA^Ala^, mt-tRNA^Thr^, and mt-tRNA^Tyr^) (**Supplementary Figure 1A**). As expected, ELAC2 was able to efficiently process nu-tRNA^Gly^ and mt-tRNA^Gln^ without additional factors and was even slightly inhibited by TRMT10C–SDR5C1 (**Figure 1C**). By contrast, processing of mt-tRNA^Ala^, mt-tRNA^Thr^, and mt-tRNA^Tyr^ was strictly dependent on the presence of TRMT10C–SDR5C1 (**Figure 1C**). The same results were obtained with mt-tRNA^Gln^ transcripts containing the 3’-trailer of pre-mt-tRNA^Tyr^, confirming that the observed differences in TRMT10C–SDR5C1 dependence are only due to the properties of the tRNAs themselves and not due to their 3’-trailer sequences (**Supplementary Figure 1B**). To confirm that the differential TRMT10C– SDR5C1-dependence of mt-tRNA processing indeed relates specifically to the degeneracy of the tRNA elbow, we generated three mutants of mt-tRNA^Gln^ containing either a G19C single mutation, a G18C/G19C double mutation, or a transplantation of the T-loop from mt-tRNA^Tyr^ (denoted T-loop^Tyr^), respectively leading to a weak, moderate, and strong disruption of the canonical elbow. While G19C has virtually no effect on 3’-processing by ELAC2 in the absence of TRMT10C– SDR5C1, G18C/G19C and T-loop^Tyr^ nearly completely abolished 3’-processing by ELAC2 alone (**Figure 1D**). Importantly, in all cases, these processing defects could be rescued by the addition of TRMT10C–SDR5C1. Taken together, these results demonstrate that the dependence of ELAC2 on additional protein factors for 3’-processing is related to the structural degeneracy of mt-tRNAs, and that TRMT10C–SDR5C1 compensates for the absence of canonical tRNA elbow structures.

### Structures of ELAC2 in complex with TRMT10C– SDR5C1 and mitochondrial tRNAs

To determine the molecular basis of tRNA recognition and 3’-processing by ELAC2, we reconstituted RNase Z with either the canonical pre-mt-tRNA^Gln^ (mt-RNase Z^Gln^) or the highly degenerated pre-mt-tRNA^Tyr^ substrate (mt-RNase Z^Tyr^) and a catalytically inactive variant of ELAC2 harboring a D550N mutation (ELAC2^mut^)^53^ **(Supplementary Figure 2)**. Mt-RNase Z^Tyr^ was reconstituted with a 5’-leaderless pre-mt-tRNA^Tyr^ substrate, while mt-RNase Z^Gln^ was reconstituted by first processing a pre-mt-tRNA^Gln^ substrate with 5’-leader and 3’-trailer in the presence of TRMT10C–SDR5C1 and PRORP, followed by the subsequent addition of ELAC2^mut^. Although TRMT10C–SDR5C1 is required only for 5’-processing but not 3’-processing of mt-tRNA^Gln^, the pre-mt-tRNA^Gln^ remained stably bound to TRMT10C– SDR5C1 in the mt-RNase Z^Gln^ complex **(Supplementary Figure 2D)**. We then determined the structures of mt-RNase Z^Gln^ and mt-RNase Z^Tyr^ using single-particle cryo-EM at global resolutions of 3.4 and 3.2 Å (Map^Gln^ and Map^Tyr^), with the ELAC2 region at 4.0 and 3.6 Å after focused refinement, respectively **(Supplementary Figure 3,4)**. From the mt-RNase Z^Tyr^ dataset, the ELAC2 core was further resolved at a higher resolution of 3.2 Å (**Supplementary Figure 3,4**). This allowed us to fit and remodel the structures of TRMT10C, the SDR5C1 tetramer, and pre-tRNA^Tyr^ based on the previously determined mt-RNase P complex structure (mt-RNase P^Tyr^) and the AlphaFold model of ELAC2^54^, resulting in complete structural models of the mt-RNase Z^Tyr^ and mt-RNase Z^Gln^ complexes **(Figure 2, Table 1)**.

**Table 1.** Cryo-EM data collection, refinement and validation statistics.

|  | Mt-RNase Z<sup>Gln</sup> complex<br>(EMDB-19455)<br>(PDB 8RR3) | Mt-RNase Z<sup>Tyr</sup> complex<br>(EMDB-19457)<br>(PDB 8RR4) |
| --- | --- | --- |
| **Data collection and processing** |  |  |
| Magnification | 105,000 x | 105,000 x |
| Acceleration Voltage (kV) | 300 | 300 |
| Electron exposure (e-/Å<sup>2</sup>) | 42.0 | 39.6 |
| Nominal defocus range (μm) | 0.5 – 2.5 | 0.5 – 2.5 |
| Pixel size (Å) | 0.834 | 0.834 |
| Symmetry imposed | C1 | C1 |
| Initial particle images (no.) | 12,168,828 | 5,987,454 |
| Final particle images (no.) | 75,812 | 28,023 |
| Map resolution (Å) | 3.4 | 3.2 |
| FSC threshold | 0.143 | 0.143 |
| Map resolution range (Å) | 9 – 2.9 | 11 – 2.8 |
| Map sharpening *B* factor (Å<sup>2</sup>) |  |  |
| Consensus map | −47.4 | −56.6 |
| Composite map | 0 | 0 |
| **Refinement** |  |  |
| Initial models (PDB) | 7ONU | 7ONU |
| Initial models (AlphaFoldDB) | Q9BQ52 | Q9BQ52 |
| Model resolution (Å) | 3.5 | 3.4 |
| FSC threshold | 0.5 | 0.5 |
| Model composition |  |  |
| Non-hydrogen atoms | 16,294 | 16,539 |
| Protein residues | 1915 | 1961 |
| Nucleotide residues | 81 | 76 |
| Ligands | Zn: 2, SAH:1 | Zn: 2, SAH:1 |
| *B* factors (Å<sup>2</sup>) |  |  |
| Protein | 30/312/130 | 33/367/154 |
| Nucleotide | 72/512/170 | 61/531/196 |
| Ligand | 275/309/292 | 356/382/369 |
| R.m.s. deviations |  |  |
| Bond lengths (Å) | 0.005 | 0.003 |
| Bond angles (°) | 1.166 | 0.602 |
| Validation |  |  |
| MolProbity score | 1.74 | 1.50 |
| Clashscore | 10.96 | 7.38 |
| Poor rotamers (%) | 0.58 | 0.50 |
| Ramachandran plot |  |  |
| Favored (%) | 96.93 | 97.52 |
| Allowed (%) | 2.97 | 2.43 |
| Disallowed (%) | 0.11 | 0.05 |

**Figure 2.**
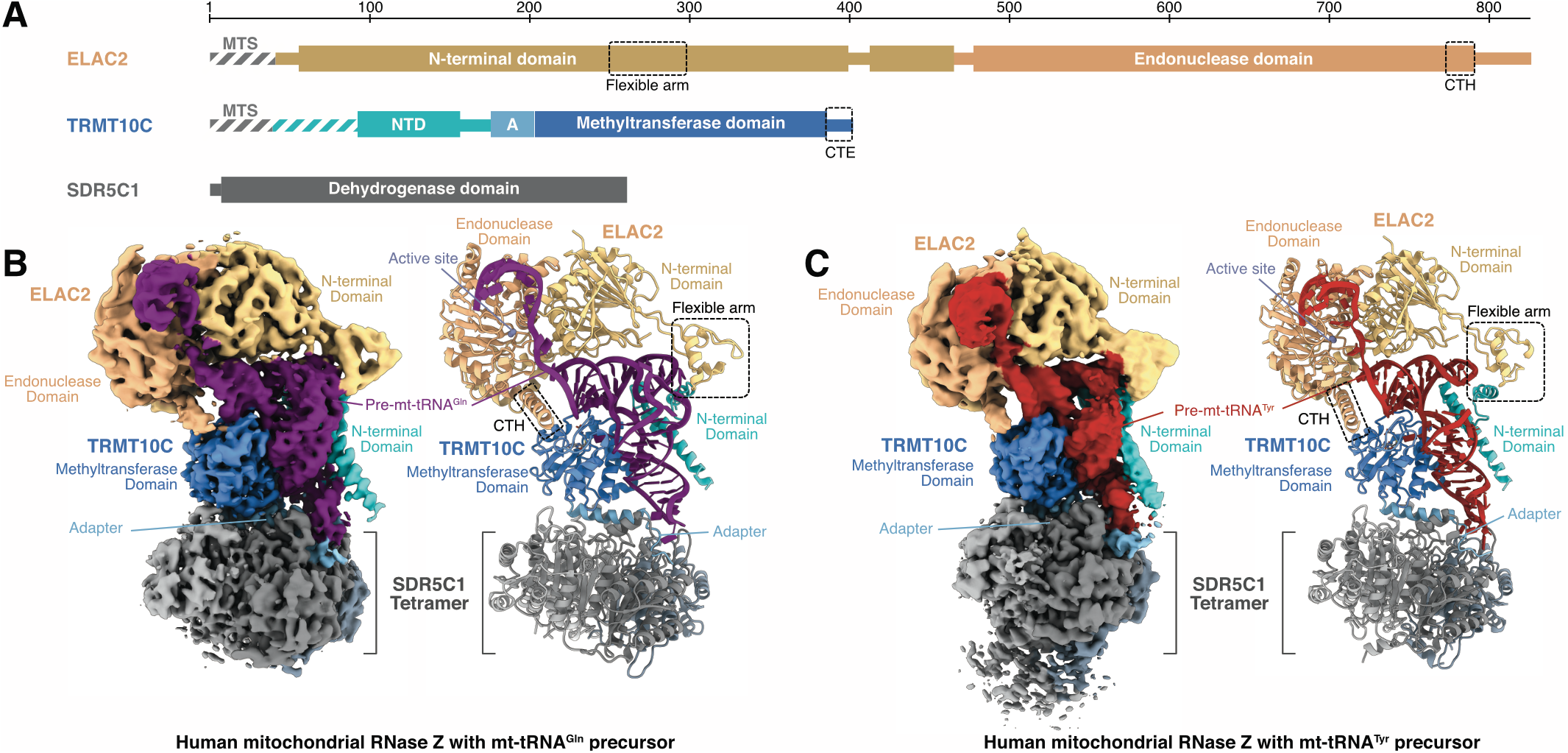
Structures of mitochondrial RNase Z complexes. **A)** Domain representation of mt-RNase Z subunits. ELAC2 domains are shown in shades of orange, TRMT10C domains shades of blue and the four SDR5C1 monomers in shades of gray throughout the manuscript. Marked by dashed boxes are the ELAC2 flexible arm and C-terminal helix (CTH), and the TRMT10C C-terminal extension (CTE). Regions of the wild-type proteins lacking in the recombinant constructs used here are marked with striped lines. Mitochondrial targeting signals (MTS) of ELAC2 and TRMT10C are indicated. **B)** Cryo-EM density map (left) and cartoon representation of the structural model (right) of the mt-RNase Z^Gln^ complex. The two catalytic Zn^2+^ ions in the active site are shown as gray spheres. The pre-mt-tRNA^Gln^ is colored in purple. ELAC2 flexible arm and CTH are marked with black dotted rectangles. **C)** Cryo-EM density map (left) and cartoon representation of the structural model (right) of mt-RNase Z^Tyr^ complex. The catalytic Zn^2+^ ions in the active site are shown as gray spheres. Pre-mt-tRNA^Tyr^ is colored in red. ELAC2 flexible arm and CTH are marked with black dotted rectangles.

Both mt-RNase Z complexes share the same architecture, which resembles that of mt-RNase P^Tyr 40^. The SDR5C1 tetramer forms a base to which TRMT10C is anchored via its “adapter helix„, and together they form a platform that binds the pre-tRNA substrate through interactions with all four tRNA subdomains. ELAC2 binds on top of the pre-tRNA and occupies the same position as PRORP in the RNase P complex, showing that binding of these two endoribonucleases is mutually exclusive^40^. ELAC2 folds into two β-lactamase domains (N-terminal domain and endonuclease domain), which form extensive interactions with the pre-tRNA via the acceptor stem, T arm and 3’-trailer. Additional contacts with the T-loop and elbow region are formed by the ∼50 residue long “flexible arm„ insertion in the ELAC2-NTD. ELAC2 also contacts TRMT10C via its flexible arm and a C-terminal helical extension. In both mt-RNase Z complexes, ELAC2 exhibits conformational variability with respect to the TRMT10C–SDR5C1–pre-tRNA subcomplex **(Supplementary Figure 5A,B)**.

The RNase Z-bound mt-tRNA^Gln^ and mt-tRNA^Tyr^ adopt similar L-shaped overall folds, but also exhibit important structural differences between the two complexes. First, mt-tRNA^Gln^ and mt-tRNA^Tyr^ adopt distinct anticodon stem-loop topologies, which are recognized by TRMT10C via different sets of interactions (**Supplementary Figure 5C,D**). Thus, the mechanism of pre-tRNA anticodon loop recognition by TRMT10C– SDR5C1 appears more versatile than previously suggested based on the mtRNase P^Tyr^ structure (**Supplementary Figure 5E,F**)^40^. Second, while pre-mt-tRNA^Gln^ adopts a stable canonical elbow structure, no stable D-loop–T-loop tertiary interactions are formed in pre-mt-tRNA^Tyr^. Consequently, the mt-RNase Z^Gln^ and mt-RNase Z^Tyr^ complexes differ with respect to their specific protein-RNA and protein-protein interactions.

### Recognition of canonical mitochondrial tRNAs by ELAC2

The structure of mt-RNase Z^Gln^ reveals how ELAC2 recognizes and interacts with canonical pre-tRNA substrates. The pre-mt-tRNA^Gln^ adopts a compact L-shaped structure, from which the 3’-trailer extends as a four nucleotides long single stranded region, followed by a short stem-loop **(Figure 3A,B)**. The T-loop of mt-tRNA^Gln^ adopts a characteristic pentanucleotide U-turn structure^55^, which establishes mutually stabilizing tertiary interactions with nucleobases G18 and G19 from the D-loop to form the canonical tRNA elbow **(Figure 3C,D)**. This includes a G19-C56 Watson-Crick base-pair at the distal end of the elbow, which represents one of the most highly conserved features of canonical tRNAs^22^. These interactions in the mt-tRNA^Gln^ elbow are stable despite a large TRMT10C-induced distortion of the D-stem of up to 10 Å compared to free tRNA^56^ **(Supplementary Figure 6A)**.

**Figure 3.**
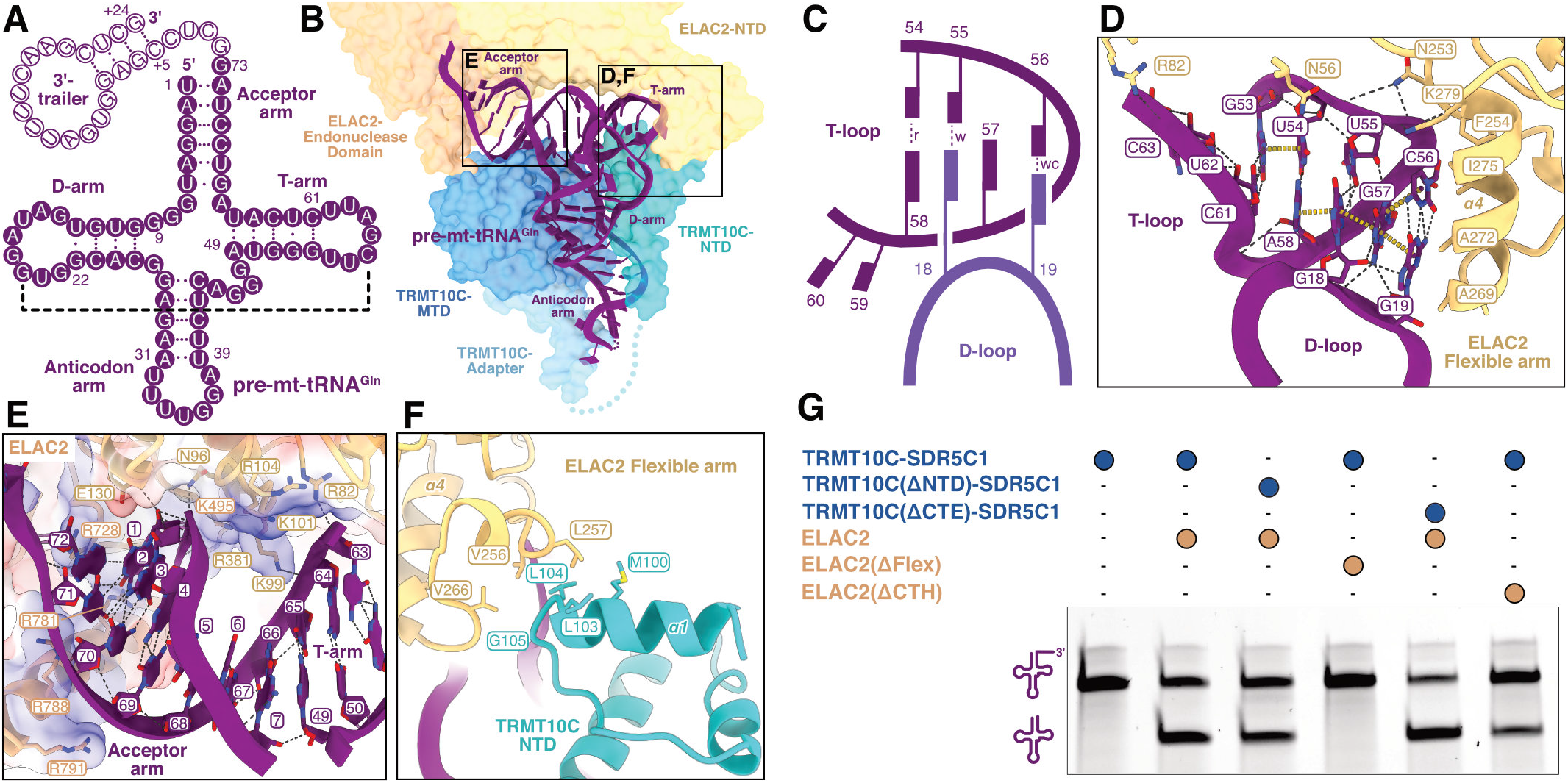
Mechanism of canonical mt-tRNA recognition by ELAC2. **A)** Secondary structure representation of pre-mt-tRNA^Gln^. The tRNA subdomains and the 3’-trailer are indicated. Nucleotides of the mature tRNA are shown as solid circles, 3’-trailer nucleotides are shown as hollow circles. G-C, A-U and G-U base pairs are marked with three, two or one dots, respectively. The tertiary G19-C56 base pair is indicated by a black dashed line. **B)** Overview of the structure and interactions of pre-mt-tRNA^Gln^ in the mt-RNase Z^Gln^ complex. ELAC2 and TRMT10C are shown as transparent surfaces and the pre-mt-tRNA^Gln^ is shown in cartoon representation. Regions shown in detail in **D-F** are indicated. **C)** Schematic representation of the tertiary interactions between the D-loop and T-loop in the mt-tRNA^Gln^ elbow. Nucleobases are represented as rectangular blocks connected to the ribose-phosphate backbone represented as solid curved lines. Watson-Crick (wc), wobble (w) or Reverse-Hoogsteen (r) base pairs are indicated as dotted lines. **D)** Recognition of the mt-tRNA^Gln^ elbow by ELAC2 flexible arm. ELAC2 flexible arm sidechains within 4.5 Å of the tRNA are shown as sticks. Hydrogen bonds and stacking interactions are shown as black and yellow dotted lines, respectively. Helix 4 (α4) of ELAC2, which forms hydrophobic interactions with the G19-C56 base-pair, is marked. **E)** Interactions of ELAC2 with the pre-mt-tRNA^Gln^ backbone in the acceptor and T arm. Surface electrostatic potential of ELAC2 is shown transparently as a three-color gradient scheme from −24 to +15 kcal mol^-1^e^-1^ (red: negative, white: neutral, blue: positive). ELAC2 sidechains within 4.5 Å of the tRNA are shown as sticks. **F)** Interactions of ELAC2 flexible arm with TRMT10C-NTD. Sidechains within 4.5 Å of the interface are shown as sticks. Helix 4 (α4) of ELAC2 and helix 1 (α1) of TRMT10C are labelled. **G)***In vitro* cleavage assays showing the effect of ELAC2 and TRMT10C deletion mutants on 3’-processing of pre-mt-tRNA^Gln^. Representative gel of three independent replicates.

ELAC2 clamps the acceptor arm–T-arm “minihelix„ of pre-tRNA^Gln^ between its N-terminal and endonuclease domains **(Figure 3B)**. The ELAC2-NTD and flexible arm form extensive contacts along the T-stem and with several conserved nucleobases of the T-loop, while the acceptor arm is inserted between the ELAC2 endonuclease domain and C-terminal helix (CTH). Both interfaces are mediated primarily by electrostatic interactions between basic protein side chains and the ribose-phosphate backbone of the Trna **(Figure 3E)**. At the distal end of the tRNA elbow, a hydrophobic patch (res. 268 to 280 in helix 4) in the globular subdomain of the ELAC2 flexible arm interacts with the conserved G19-C56 tertiary base-pair, stacking against its extended aromatic ring system **(Figure 3D)**. This interaction appears to be relatively stable, as the flexible arm shows lower conformational variability and atomic *B*-factors compared to other parts of ELAC2 **(Supplementary Figure 6B)**. This suggests that the canonical tRNA elbow structure serves as an important determinant for TRMT10C–SDR5C1-independent recognition of canonical mitochondrial tRNAs by ELAC2, consistent with our biochemical data. ELAC2 further interacts with the 3’-trailer RNA via a positively charged patch in the endonuclease domain **(Supplementary Figure 6c)**, although the resolution is not sufficient to resolve specific interactions. Together, these interactions position the 3’-end of the tRNA near the endonuclease active site of ELAC2.

Although TRMT10C–SDR5C1 is not required for efficient processing of pre-mt-tRNA^Gln^, ELAC2 also interacts with TRMT10C via two interfaces in the mt-RNase Z^Gln^ complex. The first is formed near the tRNA elbow and involves hydrophobic interactions between the ELAC2 flexible arm and the TRMT10C-NTD (**Figure 3F)**. This interface is 1.9-fold smaller than the adjacent interface between the flexible arm and the tRNA (158 Å^2^ between flexible arm and TRMT10C-NTD vs. 308 Å^2^ between flexible arm and tRNA), suggesting that the latter interface plays the primary role in ELAC2 binding to mt-tRNA^Gln^. Consistent with this, deletion of the ELAC2 flexible arm (Δ250-298) abolishes 3’-processing of mt-tRNA^Gln^, whereas deletion of the TRMT10C-NTD (Δ1-124) has no major effect **(Figure 3G)**. This shows that stabilization of the flexible arm is crucial for productive substrate binding by ELAC2, and that this is achieved primarily through interactions with the tRNA elbow rather than with the TRMT10C-NTD. The second interface is formed between the ELAC2-CTH and the TRMT10C methyltransferase domain (MTD) and C-terminal extension (CTE). This interface is only observable at low map threshold, indicating a dynamic interaction interface **(Supplementary Figure 6D)**. While deletion of the ELAC2 CTH (Δ772-826) reduces 3’-processing of mt-tRNA^Gln^, deletion of the TRMT10C CTE (Δ385-403) shows no effect **(Figure 3G)**. Thus, this interface also does not seem to play a crucial role in canonical pre-tRNA recognition and processing. Taken together, these data show that ELAC2 interacts with canonical tRNAs primarily via direct protein-RNA interactions with the acceptor stem, T-arm and elbow region, while protein-protein interactions with TRMT10C are dispensable.

### Model for nuclear tRNA recognition by ELAC2

The structure of mt-RNase Z^Gln^ also serves as a blueprint to understand the mechanism of nu-tRNA processing. The structural and biochemical results for mt-tRNA^Gln^ suggest a conserved substrate recognition mechanism for canonical mitochondrial and nuclear tRNAs **(Figure 1)**. The structure of mt-RNase Z^Gln^ thus enables us to construct a model of substrate-engaged nuclear RNase Z by superimposing human nu-tRNA^Gly^ onto the complex and omitting TRMT10C–SDR5C1 **(Figure 4A)**^57^. Structural comparison of free yeast nu-tRNA^Phe^ and mt-tRNA^Gln^ in mt-RNase Z^Gln^ complex shows that the tRNA structural elements recognized by ELAC2 remain unperturbed between free and TRMT10C–SDR5C1-bound tRNAs^56^ **(Supplementary Figure 6A)**. Consequently, all ELAC2–RNA interactions observed in mt-RNase Z^Gln^, in particular those of the flexible arm with the canonical elbow structure, can be established by a free nuclear tRNA without major structural rearrangements **(Figure 4B)**. This strongly suggests that the canonical elbow structure also serves as major determinant for nu-tRNA recognition by ELAC2. Consistent with this, mutational disruption of the elbow in nu-tRNA^Gly^ by transplantation of the T-loop from mt-tRNA^Tyr^ results in the complete loss of 3’-end processing, while the less disruptive G18C/G19C mutation does not significantly affect nu-tRNA^Gly^ processing **(Figure 4C)**. The processing defect in pre-nu-tRNA^Gly^(T-loop^Tyr^) is rescued by addition of TRMT10C–SDR5C1, demonstrating that disruption of the canonical elbow in nu-tRNAs leads to the same dependency on TRMT10C– SDR5C1 as observed for degenerated mt-tRNAs.

**Figure 4.**
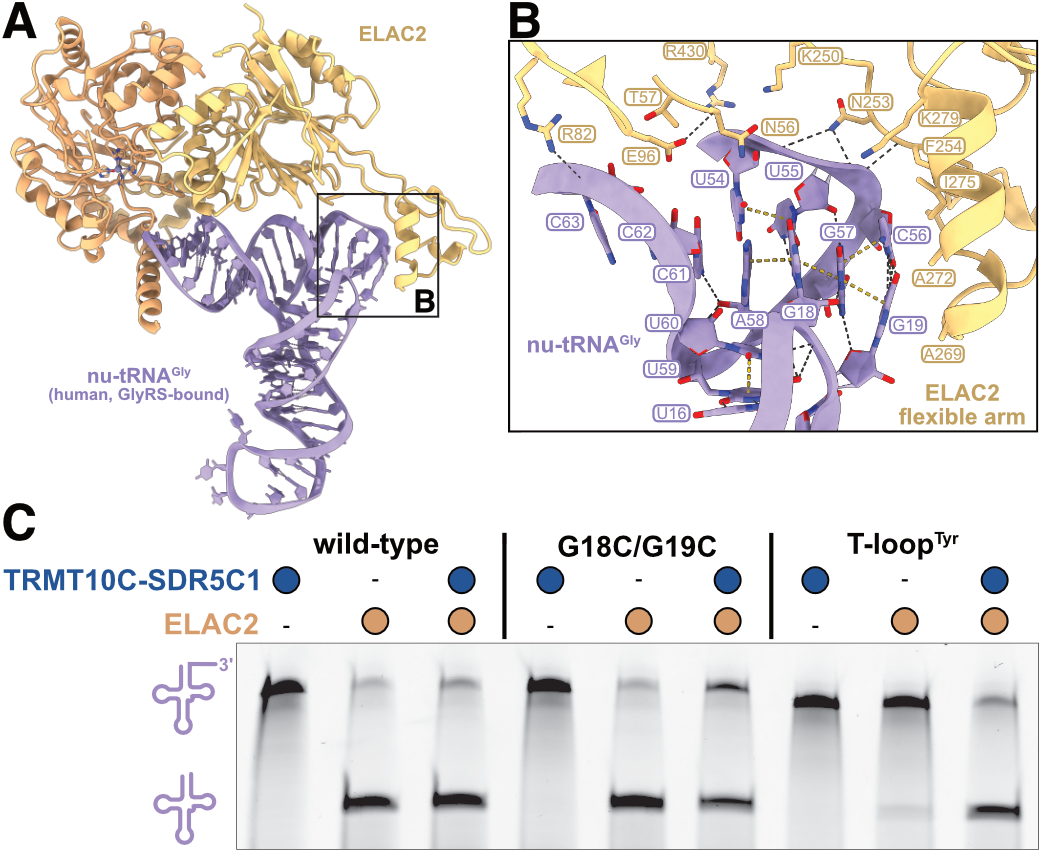
Nuclear tRNA 3’-processing by ELAC2. **A)** Model of a nuclear tRNA bound to ELAC2. Human nu-tRNA^Gly^ (bound to the glycyl-tRNA synthetase; PDB: 5E6M)^72^ (light purple) was superimposed via its acceptor and T-arms to mt-tRNA^Gln^ in the mt-RNase Z complex. TRMT10C and SDR5C1 are omitted. The region shown in **B** is indicated. **B)** Model for nu-tRNA^Gly^ elbow recognition by ELAC2 flexible arm. ELAC2 sidechains within 4.5 Å of tRNA are shown as sticks. **C)** *In vitro* cleavage assays demonstrating the critical role of the elbow region in nu-tRNA^Gly^ for 3’-processing by ELAC2. Representative gel of three independent replicates.

In summary, our data suggest how ELAC2 recognizes and interacts with canonical tRNAs both in mitochondria and the nucleus and provide a plausible model for nuclear RNase Z.

### ELAC2–TRMT10C interactions facilitate processing of degenerate mitochondrial tRNAs

The structure of mt-RNase Z^Tyr^ reveals how TRMT10C– SDR5C1 facilitate processing of structurally degenerate mt-tRNAs **(Figure 5)**. Overall, interactions between ELAC2 and pre-tRNA^Tyr^ involve largely the same ribose-phosphate interactions in the acceptor–T-arm minihelix and the 3’-trailer as observed in mt-RNase Z^Gln^ **(Figure 5A,B,E) (Supplementary Figure 7A-C)**. However, the tRNA elbow structure that is recognized by ELAC2 in canonical tRNAs is absent in mt-tRNA^Tyr^, as no stable tertiary interactions are formed between the D- and T-loops **(Figure 5C,D)**. This results in a higher structural flexibility in the elbow region of pre-mt-tRNA^Tyr^ compared to pre-mt-tRNA^Gln^, with the D-loop, and A56 and A57 at the tip of the T-loop poorly resolved in the EM density. Consequently, the extensive interactions between the ELAC2 flexible arm and the elbow observed in mt-RNase Z^Gln^ are absent in mt-RNase Z^Tyr^, resulting in a 3-fold smaller protein–RNA interface (98 Å^2^ buried surface area in mt-RNase Z^Tyr^ compared to 308 Å^2^ in mt-RNase Z^Gln^). By contrast, the interaction interface between the ELAC2 flexible arm and the TRMT10C-NTD is 1.9-fold larger in mt-RNase Z^Tyr^ than in mt-RNase Z^Gln^ (295 Å^2^ in mt-RNase Z^Tyr^ vs. 158 Å^2^ in mt-RNase Z^Gln^). This larger protein-protein interface in mt-RNase Z^Tyr^ is possible due to a reorientation of the ELAC2 flexible arm and TRMT10C-NTD towards each other, which positions the TRMT10C-NTD directly underneath the globular subdomain of the flexible arm **(Figure 5F)**. This predominantly hydrophobic interface involves M100, L103 and L104 of TRMT10C and V256, L257, K260, V266 and G267 of ELAC2, and may be further stabilized by electrostatic interactions between ELAC2 K260 and TRMT10C E96 and/or E99. As in mt-RNase Z^Gln^, a second conformationally dynamic interface is likely formed in mt-RNase Z^Tyr^ between the CTH of ELAC2 and the MTD and CTE of TRMT10C, which may involve an electrostatic interaction between ELAC2 R791 and TRMT10C D339 **(Supplementary Figure 7C)**.

**Figure 5.**
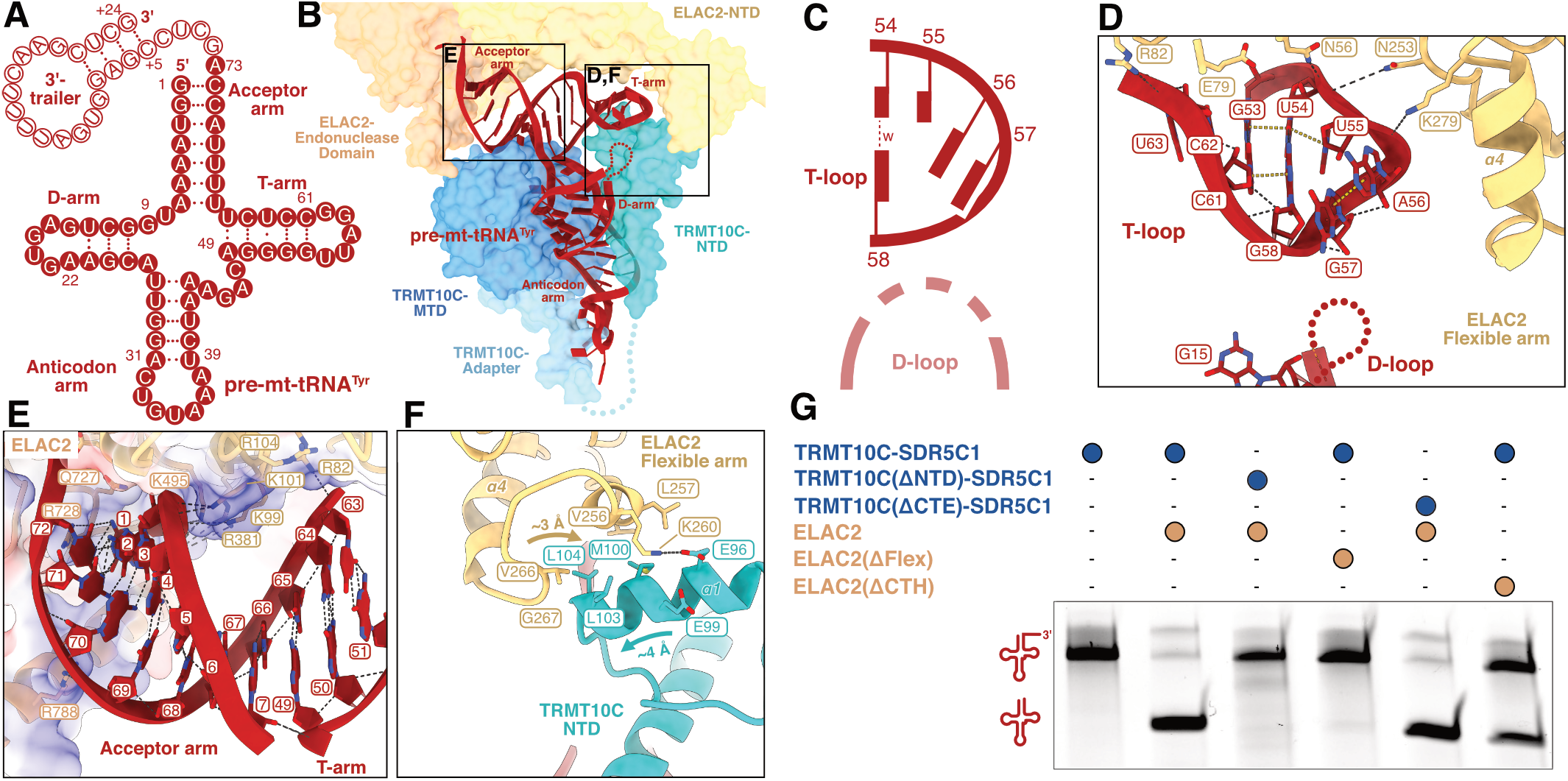
TRMT10C–SDR5C1 facilitate recognition of non-canonical mt-tRNAs by ELAC2. **A)** Secondary structure representation of pre-mt-tRNA^Tyr^. Nucleotides of the mature tRNA are shown as solid circles; the 3’-trailer is shown as open circles. **B)** Overview of the structure and interactions of pre-mt-tRNA^Tyr^ in the mt-RNase Z^Tyr^ complex. Representation as in **Fig. 3B**. Regions shown in detail in **D-F** are indicated. **C)** Schematic representation of the tertiary interactions between D-loop and T-loop in the mt-tRNA^Tyr^ elbow region. Representation as in **Fig. 3C**. **D)** Interactions between mt-tRNA^Tyr^ elbow and ELAC2 flexible arm. ELAC2 flexible arm sidechains within 4.5 Å of the tRNA are shown as sticks. **E)** Interactions of ELAC2 with the pre-mt-tRNA^Tyr^ backbone in the acceptor and T arms. Surface electrostatic potential of ELAC2 is shown transparently as a three-color gradient scheme from −18 to +17 kcal mol^-1^e^-1^ (red: negative, white: neutral, blue: positive). ELAC2 sidechains within 4.5 Å of the tRNA are shown as sticks. **F)** Interactions of ELAC2 flexible arm with TRMT10C-NTD. Sidechains within 4.5 Å of the interface are shown as sticks. Helix 4 (α4) of ELAC2 and helix 1 (α1) of TRMT10C are labelled. **G)** *In vitro* cleavage assays demonstrating the roles of ELAC2 flexible arm and TRMT10C-NTD in 3’-processing of pre-mt-tRNA^Tyr^. Representative gel of three independent replicates.

To determine the role of the two ELAC2– TRMT10C interfaces in processing of non-canonical mt-tRNA substrates, we tested deletion mutants of each of the involved domains in our *in vitro* cleavage assay **(Figure 5G)**. The most severe effect was observed for the ELAC2 flexible arm deletion, which abolished 3’-processing of pre-mt-tRNA^Tyr^, similar to pre-mt-tRNA^Gln^ **(Figure 3G, 5G)**. However, in contrast to pre-mt-tRNA^Gln^, deletion of the NTD of TRMT10C also abolishes TRMT10C–SDR5C1-supported 3’-processing of pre-mt-tRNA^Tyr^. This demonstrates that the TRMT10C-NTD, while dispensable on canonical tRNAs, is critical for TRMT10C–SDR5C1-dependent 3’-processing by ELAC2 on degenerated mt-tRNA substrates. A less pronounced activity loss was observed for the ELAC2-CTH deletion. Interestingly, this effect is exacerbated on mt-tRNA^Gln^ variants with mutationally disrupted elbow region **(Supplementary Figure 7D)**, suggesting that the CTH may facilitate processing of destabilized tRNAs by tethering ELAC2 to the TRMT10C–SDR5C1–pre-tRNA complex. This effect is likely mediated by direct interactions of the ELAC2-CTH with the tRNA acceptor arm and TRMT10C-MTD, but not with the TRMT10C-CTE, as deletion of the latter has no negative effect on ELAC2-mediated 3’-cleavage **(Figure 5G)**.

Taken together, our structural and biochemical data demonstrate that protein-protein interactions between ELAC2 and TRMT10C compensate for the lack of extensive protein-RNA interactions with otherwise conserved structural elements in degenerate mt-tRNAs.

### Active site organization of ELAC2

The active site organization in human ELAC2 is similar to that of yeast RNase Z_L_ (Sce-RNase Z_L_) or *Bacillus subitilis* (Bsu) RNase Z^44,58^. In both mt-RNase Z complexes, the active site contains clear density for two putative zinc ions **(Supplementary Figure 8A,B)**. Zn^2+^_A_ is coordinated by H546, H548, H644 and D666, while Zn^2+^_B_ is coordinated by H551, D666 and H724. The residue mutated in our catalytically inactive variant of ELAC2, D550N, lies adjacent to the active site. Although N550 does not coordinate Zn^2+^_B_ in our structures, D550 likely forms part of the coordination sphere of Zn^2+^_B_ in wild-type ELAC2. However, structural comparisons to Sce-RNase Z_L_ and Bsu-RNase Z show that the D550N mutation does not significantly alter the active site architecture or relative positioning of catalytic residues and ions (RMSD at active site: Sce-RNase Z_L_ – 0.84 Å, Bsu-RNase Z – 0.63 Å) **(Supplementary Figure 8C)**.

Due to the large conformational variability of ELAC2 with respect to TRMT10C–SDR5C1–pre-tRNA, the pre-tRNA adopts an ensemble of “productive„ and “non-productive„ conformations near the 3’ cleavage site. To better resolve the “productive„ active site conformation, we performed a 3-D variability analysis of mt-RNase Z^Tyr^ particles with respect to the ELAC2 conformational state **(Supplementary Figure 8D)**. This resulted in a class where the pre-mt-tRNA^Tyr^ adopts a “contracted„ conformation allowing the RNA to be positioned in the ELAC2 active site. Although the resulting maps are of limited resolution (∼4.1 Å), comparison with the pre-tRNA-bound structure of the Bsu-RNase Z homodimer allowed us place the RNA in the active site of ELAC2 such that the scissile phosphodiester is positioned next to the Zn^2+^ ions^44,59^ **(Supplementary Figure 8E,F)**. This model shows that D550, proposed to act as the general base catalyst^44,60^, and H702, proposed to protonate the leaving group^60^, are positioned near the scissile phosphate. The discriminator nucleotide (N73) is likely stabilized by K700, while the nucleobase in position 75 is stabilized in a groove formed by residues L547, H548, C645, K646, H647 and N583. Both interactions may contribute to positioning the scissile phosphodiester bond in the active site.

In summary, our structural data provide a pre-catalytic model of substrate-engaged ELAC2, which suggests a conserved catalytic mechanism of RNase Z enzymes.

### Structural basis for sequential tRNA processing in mitochondria and nucleus

For most human mitochondrial and nuclear tRNAs, maturation proceeds sequentially, starting with 5’-end processing by RNase P followed by 3’-end processing by ELAC2^42,61,62^. Superimposing mt- or nu-RNase P complexes with RNase Z reveals steric clashes between PRORP or nu-RNase P and ELAC2. This suggests that association of 5’- and 3’-processing enzymes to the pre-tRNA is mutually exclusive **(Supplementary Figure 9A,B)**. Furthermore, the 5’-phosphate of the ELAC2-bound pre-tRNA is displaced by ∼6 Å compared to mt-RNase P-bound state^40^, and is buried near the ELAC2 core such that binding of a 5’-unprocessed pre-tRNA would be disfavored due to steric clashes between the 5’-leader and ELAC2. In particular, a loop between residues 726 and 731, and the CTH of ELAC2 would clash with the 5’-leader at positions −2 and −3, respectively **(Supplementary Figure 9C,D)**. Deletion of the ELAC2-CTH alone does not abolish the discrimination against pre-tRNA with a 5’-leader by ELAC2, suggesting that multiple elements of ELAC2 are involved in ensuring hierarchical processing **(Supplementary Figure 9E)**.

In conclusion, we find that sequential tRNA 5’- and 3’-end processing requires an exchange of RNase P and RNase Z enzymes, and this strict processing order is likely ensured by steric discrimination against 5’-unprocessed pre-tRNAs by ELAC2.

## DISCUSSION

Here we present structures of human mt-RNase Z bound to two structurally divergent pre-tRNA substrates, which serve as models for both mitochondrial and nuclear tRNA 3’-processing. Together with biochemical data, the structures reveal the molecular determinants for substrate recognition by RNase Z and explain why ELAC2 requires TRMT10C–SDR5C1 to process most mt-tRNAs. Furthermore, they provide insight into the active site arrangement of RNase Z and enable us to propose a molecular model for sequential processing of nuclear and mitochondrial tRNAs. Taken together, our results elucidate the molecular mechanism of tRNA 3’-end processing of human mitochondrial and nuclear tRNAs by a single RNase Z enzyme.

Most tRNAs encoded in human mitochondria are highly degenerate, and differ significantly in sequence, structure, and stability from canonical tRNAs encoded in the nucleus^28,63^. Most notably, mt-tRNAs often lack universally conserved tRNA structural features that are used by many tRNA-processing factors to recognize nu-tRNAs^23,28,35,40^. ELAC2 catalyzes tRNA 3’-processing both in the nucleus and mitochondria ^31,42,49^, and thus must recognize pre-tRNAs from both compartments. Our structural and biochemical data reveal that ELAC2 recognizes canonical nuclear and degenerated mitochondrial tRNAs by distinct mechanisms.

For canonical tRNAs, the elbow structure is recognized by the flexible arm of ELAC2 through interactions with the conserved T-loop and G19-C56 pair. The flexible arm and endonuclease domain of ELAC2 clamp the tRNA core, which positions the 3’ cleavage site near the active site. Hence, disruption of the tRNA elbow would impair the stabilization of the ELAC2 flexible arm and disfavor productive positioning of the 3’ cleavage site. All protein-RNA interactions observed in mt-RNase Z^Gln^ can be formed also in absence of TRMT10C– SDR5C1. These protein-RNA interactions observed in mt-RNase Z^Gln^ are likely the major determinants for recognizing canonical pre-tRNA substrates both in mitochondria and the nucleus. This is also similar to the mechanism of substrate binding by the *B. subtilis* RNase Z_S_ homodimer, suggesting a conserved mechanism of canonical tRNA recognition between long- and short-form RNase Zs^44^.

Most human mt-tRNAs contain reduced or variable D- and T-loops that cannot form the canonical elbow structure^28,29,64,65^. Our analysis shows that TRMT10C–SDR5C1 specifically enables 3’-end processing of mt-tRNAs in which the canonical elbow is degenerated. The structure of mt-RNase Z^Tyr^ shows that TRMT10C–SDR5C1 facilitates processing of such mt-tRNAs by stabilizing ELAC2 via protein-protein interactions with TRMT10C, which compensate for the loss of interactions between ELAC2 and the tRNA elbow. Thus, the TRMT10C–SDR5C1 complex appears to play a twofold role in mt-RNase Z: on the one hand, it recognizes and stabilizes intrinsically flexible mt-tRNAs in a common L-shaped fold; on the other, it acts as a “prosthetic„ extension for degenerated mt-tRNAs lacking stable elbow structures and provides compensatory anchor points for ELAC2. This compensatory role of the TRMT10C–SDR5C1 complex is analogous to its function in mt-RNase P, where it similarly stabilizes the mt-tRNA substrate and mediates binding of the PRORP subunit for 5’-processing^40,66^.

The structural data also provide insight into the catalytic mechanism of ELAC2. In the catalytically “productive„ conformation, the phosphodiester group of nucleotide 74 is positioned in the ELAC2 active site, between to the two Zn^2+^ ions, the putative general base D550, and H702, which was proposed to stabilize the leaving group^60^. The resulting pre-catalytic configuration of ELAC2 is very similar to that previously proposed for *Bsu*-RNase Z, suggesting a conserved catalytic mechanism among RNase Zs^44,47,60^.

Moreover, our results support the previously proposed role of TRMT10C–SDR5C1 as a mitochondrial tRNA maturation platform^41^. As the structures of mt-RNase Z and mt-RNase P show, TRMT10C requires neither PRORP nor ELAC2 to bind the pre-tRNA, and none of the extensive TRMT10C–tRNA interactions are affected by either 5’- or 3’-processing^66^. Consistent with this, our reconstitution of the mt-RNase Z^Gln^ complex demonstrates that 5’-processed pre-tRNA^Gln^ remains associated with TRMT10C–SDR5C1 following PRORP dissociation and during subsequent 3’-processing, despite the slightly inhibitory effect TRMT10C–SDR5C1 appears to have on ELAC2 activity on this tRNA **(Figure 1C)**. Thus, our results suggest that mt-tRNAs remain bound to the TRMT10C–SDR5C1 complex for multiple maturation steps, consistent with previous biochemical observations, as well as with structures of mt-RNaseZ^His^ and TRMT10C–SDR5C1–mt-tRNA^Ile^–TRNT1 complexes that were reported in a preprint during the preparation of this manuscript^41,67^. Additional maturation steps supported by the TRMT10C–SDR5C1 platform following 5’- and 3’-processing may include 3’-CCA-addition by TRNT1 as well as further modifications by modifying enzymes and aminoacyl tRNA synthetases (AARSs).

These observations lead us to hypothesize that the TRMT10C–SDR5C1 complex may have evolved as a common solution for multiple mt-tRNA-binding factors to the common challenge of substrate recognition posed by the structural degeneration of mt-tRNAs^33,35^. The canonical tRNA elbow structure plays a key role in cytosolic gene expression machineries, as it is recognized by many tRNA-binding factors including RNase Ps, RNase Zs, CCA-nucleotidyltransferases, AARSs, and the ribosome^23,39,47,68–70^. Its loss in mt-tRNAs thus imposes strong pressure to evolve compensatory mechanisms to maintain these functional interactions. In human mt-RNase P and mt-RNase Z, the PRORP-PPR domain and ELAC2 flexible arm, both of which classically interact with the tRNA elbow, interact with TRMT10C-NTD via the same surface on its globular subdomain. Notably, this globular subdomain first appears and becomes fixed in TRMT10 homologues of bilaterians, coinciding with the widespread erosion of mt-tRNAs^30^ **(Supplementary Figure 10)**. Thus, the TRMT10C– SDR5C1 complex may have evolved to specifically compensate for the structural erosion of mt-tRNAs and to maintain their indispensable function in mitochondrial gene expression.

In conjunction with previous data, our data allow us to propose a model for the sequential maturation of nuclear and mitochondrial tRNAs (**Figure 6**). First, RNase P binds to the pre-tRNA containing 5’-leader and 3’-trailer to catalyze 5’-processing. ELAC2 is prevented from productive binding to the same substrate at this stage due to steric discrimination against 5’-unprocessed pre-tRNAs. Following 5’-end processing, RNase P must dissociate before ELAC2 can be recruited to the pre-tRNA through interactions of the ELAC2 flexible arm with either the tRNA elbow or the TRMT10C-NTD, leading to 3’-processing. The resulting 5’- and 3’-processed tRNA can then be further matured by TRNT1, which catalyzes addition of the 3’-CCA end. In the nucleus, the enzymes carrying out these maturation steps are self-sufficient, with nuclear RNase Z acting as a single-subunit enzyme (**Figure 6A**). By contrast, in mitochondria all three maturation steps take place on pre-tRNAs stably associated with TRMT10C–SDR5C1 (**Figure 6B**).

**Figure 6.**
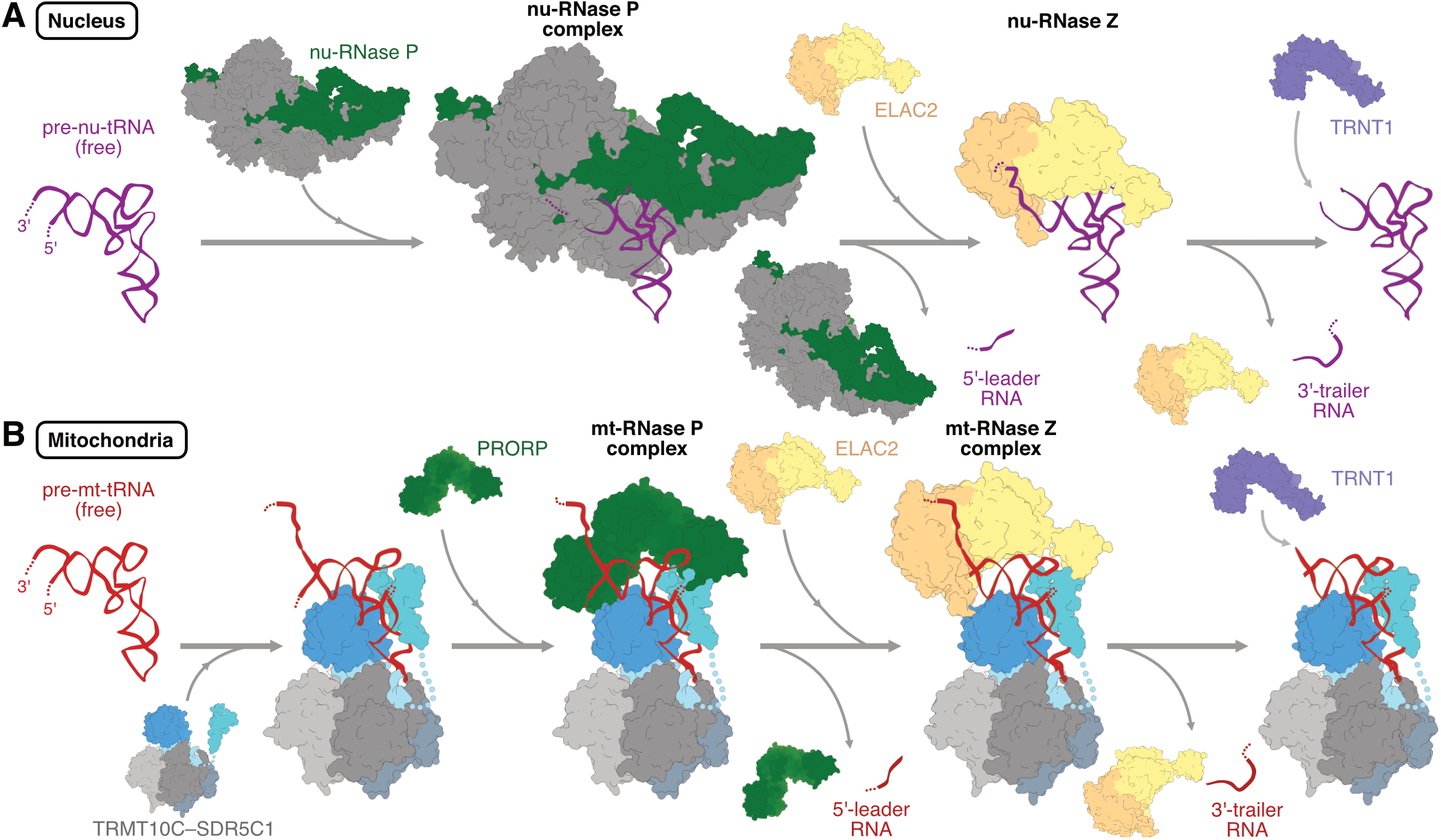
Models of nuclear and mitochondrial tRNA processing. **A)** Model of sequential tRNA maturation in the nucleus. Nuclear pre-tRNA is shown in purple; the RNA subunit of nu-RNase P is shown in green and the accessory protein subunits are shown in gray; ELAC2 is shown in shades of orange and TRNT1 in light purple. **B)** Model of sequential tRNA maturation in mitochondria. Mitochondrial pre-tRNA is shown in red; TRMT10C in shades of blue; SDR5C1 tetramer in shades of gray; PRORP in green; ELAC2 is shown in shades of orange and TRNT1 in light purple.

In summary, our results explain how ELAC2 recognizes and catalyzes 3’-end processing of two structurally divergent sets of tRNAs found in human mitochondria and the nucleus. Together with previous observations, these results further highlight the role of the conserved tRNA elbow structure in tRNA recognition and provide insight into the evolutionary history of TRMT10C–SDR5C1 as a mitochondrial tRNA maturation platform.

## METHODS

No statistical methods were used to predetermine sample size. The experiments were not randomized, and the investigators were not blinded to allocation during experiments and outcome assessment.

### Protein cloning, expression & purification

PRORP, TRMT10C and SDR5C1 were cloned, expressed and purified as previously reported^40^. Mutant variants of TRMT10C were generated by restriction-free cloning PCR using the vector encoding wildtype TRMT10C and SDR5C1 as the template. TRMT10C– SDR5C1 complexes containing mutant TRMT10C were expressed and purified following the same protocol as for the wildtype proteins. The sequence encoding ELAC2 lacking the N-terminal mitochondrial targeting sequence (Δ1–31) was PCR amplified from human cDNA obtained from B-lymphocytes. It was cloned into 438-B vector (kind gift from Scott Gradia; Addgene plasmid # 55219; http://n2t.net/addgene:55219; RRID:Addgene_55219^82^) in frame with an N-terminal 6xHis tag followed by a TEV cleavage site, followed by generation of baculovirus. ELAC2 mutants were generated by site-directed mutagenesis using Round-the-Horn PCR amplification of wild-type ELAC2 in 438-B vector. ELAC2 and its variants were expressed in an insect cell expression system using Hi-Five cells as previously described^73,83^. All subsequent steps were carried out at 4 °C. Cells were harvested by centrifugation at 238 g for 30 minutes and resuspended in lysis buffer at pH 7.5 containing 20 mM Na-HEPES, 300 mM NaCl, 40 mM imidazole, 1 mM dithiothreitol (DTT), 10% glycerol and 1x protease inhibitor cocktail (Roche) or 0.284 µg mL^−1^ leupeptin, 1.37 µg mL^−1^ pepstatin, 0.17 mg mL^−1^ phenylmethylsulfonyl fluoride and 0.33 mg ml^−1^ benzamidine. Cells were lysed by sonification, followed by sequential centrifugation in an A27 rotor (Thermo Fisher) at 26,195 xg for 30 minutes, and ultracentrifugation in a Type 45 Ti rotor (Beckman Coulter) for 60 minutes. The supernatant was filtered through filtration membranes with sieve width of 5 µm followed by 0.8 µm and applied to a HisTrap HP 5 ml column (Cytiva) equilibrated with lysis buffer. The column was washed with 10 column volumes (CV) of lysis buffer, followed by 10 CV of high-salt wash buffer (50 mM Na-HEPES, 1 M NaCl, 40 mM imidazole, 10% glycerol, 2 mM DTT, pH 7.5), followed again by 5 CV of lysis buffer. Bound proteins were eluted with 9.5 CV of elution buffer (17 mM Na-HEPES, 255 mM NaCl, 334 mM imidazole, 8.5% glycerol and 1.7 mM DTT). To the eluted protein, 1.5 mg of recombinant TEV protease (homemade) was added, followed by overnight dialysis in 50 mM Na-HEPES, 300 mM NaCl, 10% glycerol, 2 mM DTT at pH 7.5. Dialyzed eluate was reapplied to the HisTrap HP column equilibrated with buffer containing 50 mM Na-HEPES, 300 mM NaCl, 10 mM imidazole, 10% glycerol, 2 mM DTT at pH 7.5 and washed with 10 CV of the same buffer. The flow-through and wash were collected and applied to a HiTrap Heparin 5 mL column (Cytiva) equilibrated with 50 mM Na-HEPES, 150 mM NaCl, 10% glycerol and 2 mM DTT at pH 7.5. Bound proteins were eluted with a gradient of 150 to 1000 mM NaCl with 50 mM Na-HEPES, 10% glycerol and 2 mM DTT at pH 7.5. Fractions containing ELAC2 according to SDS-PAGE were pooled and further purified using a Superdex 200 Increase 10/300 GL column (Cytiva) equilibrated with 20 mM Na-HEPES, 150 mM NaCl, 10% glycerol and 5 mM DTT at pH 7.5. Fractions containing ELAC2 were concentrated using Amicon Ultra-4 30K Centrifugal Filter Devices (Merck Millipore), aliquoted, flash-frozen in liquid nitrogen and stored at −70 °C.

### Preparation of substrate RNAs

Sequences encoding the pre-tRNA substrates under the control of a T7 RNA polymerase promoter were either purchased as gBlocks (Integrated DNA Technologies) or were cloned into a pUC19 vector. Mutations were introduced by site-directed mutagenesis PCR. The initial templates were amplified by PCR using forward and reverse primers complementary to T7 promoter and the 3’ trailer sequence, respectively. PCR-amplified templates were purified using QIAquick or MinElute PCR Purification Kits (Qiagen) and used for run-off *in-vitro* transcription (IVT).

The RNA substrates for structural studies – tRNA^Tyr^ with no 5’ leader and 24 nt-long 3’ trailer (0– tRNA^Tyr^–24), and tRNA^Gln^ with 5 nt-long 5’ leader and same 24 nt-long 3’ trailer as tRNA^Tyr^ (5–tRNA^Gln^–24) – were transcribed in reactions containing 1x T7 RNA polymerase reaction buffer (NEB), 0.001% (w/v) Triton-X 100, 30 mM MgCl_2_, 4 mM NTPs, 5 U µL^-1^ T7 RNA polymerase (NEB) and 0.2 µg µL^-1^ template DNA. The Mg_2_P_2_O_7_ precipitate formed during IVT was solubilized with EDTA at a final concentration of 37 mM and removed by centrifugation. RNAs were then purified by anion exchange chromatography using a RESOURCE Q 6 mL column (Cytiva). The column was equilibrated with 9.5 CV of buffer A containing 50 mM NaCH_3_COO and 2 mM MgCl_2_, followed by application of the IVT reaction. Bound RNAs were eluted with a linear gradient from 0-100% buffer B containing 1000 mM NaCl, 50 mM NaCH_3_COO and 2 mM MgCl_2_. Elution fractions were analyzed by Urea PAGE and fractions of interest were pooled, and RNA was precipitated with NaCH_3_COO and ethanol at final concentrations of 300 mM and 70 %, respectively. Pure RNAs were dissolved in nuclease-free water and stored at –20 °C until further use.

All tRNA substrates used for biochemical analysis were transcribed in reactions containing 1x T7 RNA polymerase reaction buffer (Thermo Fisher), 2 Mm DTT, 30 mM MgCl_2_ , 6 mM of each NTP, 0.002 U µL^-1^ *E. coli* PPIase (NEB), 5 U µL^-1^ T7 RNA polymerase (Thermo Fisher) and 50 ng µL^-1^ template DNA. After incubation at 37°C for 10-14 hours, reactions were stopped by addition of an equal volume of 2x TBE-Urea Sample Buffer (Thermo Fisher) and boiling at 95 °C for 5 min. Samples were then separated by 10% Urea PAGE, visualized by UV-shadowing and bands containing the RNA of interest were cut and transferred to an RNase-free Eppendorf tube. Gel slices were crushed and RNAs extracted in buffer containing 300 mM NaCH_3_COO, 1 mM EDTA, and 20 mM Tris, pH 5. Extracted RNAs were ethanol-precipitated and stored as described above.

### RNA cleavage assays

Pre-tRNA cleavage assays were carried out in buffer containing 20 mM HEPES/KOH pH 7.4, 150 mM KCl, 3 mM MgCl_2_, 10 µM ZnCl_2_, 2 mM DTT, and 100 µM S-adenosyl-L-methionine (SAM). Reactions were set up by adding 200 nM pre-tRNA to the reaction buffer, followed by incubation for 10 min at 30 °C in the presence or absence of 800 nM TRMT10C–SDR5C1 complex. Reactions were started by the addition of 50 nM ELAC2, followed by incubation at 30 °C for 20 min, unless stated otherwise. For control reactions on 5’-leader-containing pre-tRNA substrates, 50 nM PRORP were added together with ELAC2. Reactions were stopped by addition of 2x TBE-Urea Sample Buffer (Thermo Fisher), supplemented with Proteinase K (Thermo Fisher), and incubated at 50 °C for 30 min. Samples were boiled for 5 min at 95 °C and then loaded onto a 15% TBE-Urea PAGE. Gels were subsequently soaked in TBE buffer containing 1x SYBR Gold Nucleic Acid Gel Stain (Thermo Fisher) and visualized on a Typhoon imager (Cytiva).

### Cryo-EM sample preparation and data collection

Δ1-91 TRMT10C was used for the structural studies to avoid excessive particle clustering observed with full-length TRMT10C. For assembly of mt-RNase Z^Gln^ complex, 2.7 nmol of 5–tRNA^Gln^–24 substrate was first subjected to 5’ processing with 2.7 nmol of Δ1-91 TRMT10C–SDR5C1 complex and 270 pmol of Δ1-45 PRORP in a reaction buffer at pH 8.0 containing 25 mM Tris-HCl, 150 mM NaCl, 5 mM MgCl_2_, 5 mM DTT and 200 µM SAM, and incubated at 30 °C for 16 h. The reaction buffer was then exchanged to a buffer at pH 8.0 containing 25 mM Tris-HCl, 20 mM NaCl, 20 µM ZnCl_2_, 40 µM SAM and 4 mM DTT using Zeba™ Spin Desalting Columns (Thermo Fisher). 7.0 nmol of Δ1-31 ELAC2(D550N) was added and the reaction incubated on ice for 60 minutes. For assembly of mt-RNase Z^Tyr^ complex, 2.6 nmol of 0–tRNA^Tyr^–24 substrate was mixed with 1.1 nmol of Δ1-91 TRMT10C–SDR5C1 and 3 nmol of ELAC2(D550N) in buffer at pH 8.0 containing 25 Mm Tris-HCl, 25 mM NaCl, 2 mM DTT, 10 µM ZnCl_2_ and 20 µM S-adenosyl homocysteine (SAH), and incubated on ice for 60 minutes at 4 °C. Both assembled complexes were then used for GraFix^84^ in 10 to 30% sucrose density gradient at pH 7.5 containing 25 mM Na-HEPES, 20 mM NaCl, 20 µM ZnCl_2_ and 2 mM DTT with and without a glutaraldehyde gradient (0 to 0.015% for mt-RNase Z^Gln^ and 0 to 0.025 % for mt-RNase Z^Tyr^). Ultracentrifugation was carried out in a SW 60 Ti Swinging-Bucket Rotor (Beckman Coulter) at 40,000 rpm at 10 °C for 16 h. The gradient solutions were divided into 200 uL fractions and analyzed via SDS PAGE and Nanodrop One spectrophotometer (Thermo Fisher). Fractions 14 to 17 from the gradient with glutaraldehyde crosslinker were pooled for both complexes for cryo-EM, based on SDS PAGE profiles for both samples without the crosslinker **(Supplementary Figure 2C-F)**. Pooled fractions were buffer-exchanged to 25 mM Na-HEPES, 20 mM NaCl, 20 µM ZnCl_2_ and 2 mM DTT, and concentrated using Amicon Ultra 0.5 mL 10 KDa cutoff centrifugation devices (Merck Millipore). 4 uL of the concentrated sample was applied to freshly glow-discharged R2/1 holey carbon grids (Quantifoil), blotted with blot force of 5 for 5 s using a Vitrobot Mark IV (Thermo Fisher) at 95% humidity and 4 °C, and plunge-frozen in liquid ethane.

Cryo-EM data collection was performed with SerialEM^85^ using a Titan Krios transmission electron microscope (Thermo Fisher) operated at 300 keV. Images were acquired in EFTEM mode using a GIF quantum energy filter set to a slit width of 20 eV and a K3 direct electron detector (Gatan) at a nominal magnification of 105,000 x corresponding to a calibrated pixel size of 0.834 Å pixel^-1^. Exposures were saved as non-super-resolution counting image stacks of 40 movie frames, with electron doses of 0.99 to 1.05 e^-^Å^-2^ per frame. Image stacks were acquired with stage movement per hole for the mt-RNase Z^Tyr^ dataset, and with stage movement per 3 by 3 holes with active beam-tilt compensation for the mt-RNase Z^Gln^ dataset, as implemented in SerialEM.

### Cryo-EM data processing and analysis

Image stacks were preprocessed on-the-fly with gain correction, motion correction, CTF estimation, particle picking and extraction at 2.5 Å pixel^-1^ using Warp v1.0.9^77^.

For mt-RNase Z^Gln^ dataset, 12,168,828 particles autopicked from 39,387 micrographs were subjected to 2-D classification in cryoSPARC **(Supplementary Figure 3A)**. Particles belonging to classes clearly lacking protein-like features were discarded, while the 11,090,926 particles belonging to remaining classes were divided into “good„ or “bad„ subsets. The “bad„ particle subset was used for *ab-initio* reconstruction resulting in five “bad„ references. A single “good„ reference was obtained from a previous smaller dataset of mt-RNase Z^Gln^ complex processed in cryoSPARC v4.2.1 (Structura Biotechnology)^76^. All 11,090,926 particles were subjected to supervised 3-D classification (heterogeneous refinement algorithm in cryoSPARC) using the “good„ and “bad„ initial references. 4,192,922 particles belonging to the “good„ class were subjected to another 2-D classification in cryoSPARC, from which 3,255,655 particles were selected. These particles were further subjected to unsupervised 3-D classification in RELION v3.1.0^75^, which yielded in a single class of 509,418 particles leading to an isotropic reconstruction of the TRMT10C–SDR5C1–pre-tRNA^Gln^ module. These particles were re-extracted at 0.834 Å pixel^-1^ and used for a consensus refinement and subsequent CTF refinement in RELION. To classify with respect to heterogeneity around the ELAC2-binding site, focused 3-D classification without image alignment was carried out in cryoSPARC with a mask around the ELAC2-binding site, resulting in 152,762 particles with robust ELAC2 density. After consensus refinement and a focused refinement in cryoSPARC centered on the ELAC2 density, another focused 3-D classification was carried out in cryoSPARC with mask around the 3’ trailer RNA and ELAC2 active site, resulting in a class of 75,812 particles with ordered 3’ trailer density. These particles were subjected to a consensus 3-D refinement and focused refinement in cryoSPARC centered of the ELAC2 density. The resulting consensus and ELAC2-focused maps were resharpened with a *B*-factor of 0 Å^2^ to avoid over-sharpening of the flexible regions and combined using PHENIX ^78^, resulting in a composite map of the mt-RNase Z^Gln^ complex^78^.

For mt-RNase Z^Tyr^ dataset, 5,987,454 particles autopicked from 15,344 micrographs were subjected to 2-D classification, from which particles clearly lacking RNase Z-like features were discarded **(Supplementary Figure 3B)**. The remaining 5,275,530 particles were divided into “good„ and “bad„ subsets, and the particles from the “bad„ subset were used to generate four “bad„ initial references using the *ab-initio* reconstruction algorithm in cryoSPARC. A low-resolution consensus map of mt-RNase Z^Tyr^ complex from a previous smaller dataset was used as the “good„ initial reference. All 5,275,530 particles were subjected to a supervised 3-D classification in cryoSPARC with the four “bad„ and a single “good„ initial reference. The resulting 1,780,577 particles belonging to the “good„ class were subjected to a focused 3-D classification with respect to the density around the ELAC2-binding site in RELION. 482,446 particles containing robust density at the ELAC2-binding site were re-extracted at 0.834 Å pixel^-^1, and subjected to an unsupervised 3-D classification in RELION, which yielded a single class of 227,594 particles resulting in an isotropic reconstruction of the complex. These particles were further cleaned up using 2-D classification in cryoSPARC, and a second focused 3-D classification around ELAC2 density, resulting in a class of 57,585 particles which resulted in the high-resolution map of ELAC2 at 3.2 Å. These particles were further classified with respect to the density around the ELAC2 flexible arm, resulting in a class with 28,023 particles showing ordered ELAC2 flexible arm density. These particles were subjected to a consensus refinement and a focused 3-D refinement around ELAC2 density using cryoSPARC. The resulting maps were resharpened with a *B*-factor of 0 Å^2^ to avoid over-sharpening the flexible regions and combined in PHENIX^78^, resulting in the composite map of the mt-RNase Z^Tyr^ complex.

3-D variability analysis for both complexes was performed in cryoSPARC^76^ using alignments and reference volumes from the consensus refinement, solving for two principal modes of covariance. To resolve the productive active site conformation in mt-RNase Z^Tyr^, outputs of the consensus refinement of the 57,585 particles resulting in the high-resolution map of ELAC2 were used. Particles were binned into three subsets based on their reaction coordinates along the first principal component. The subsets were independently refined with a refinement mask around the ELAC2 density, resulting in density maps for the three subsets, showing the “extended„, “relaxed„ and “contracted„ conformation of the pre-tRNA. Local resolution estimations were calculated in cryoSPARC^76^. Angular distribution plots were generated using a script packaged with Warp 1.0.9^77^.

### Model building and refinement

The initial models for TRMT10C, SDR5C1 and pre-tRNA^Tyr^ were obtained from the model of mt-RNase P^Tyr^ complex (PDB: 7ONU)^40^, and for ELAC2 from AlphaFoldDB (Accession no. Q9BQ52). The models were rigid-body-fit into the final maps of mt-RNase Z^Gln^ or mt-RNase Z^Tyr^ complex using UCSF ChimeraX 1.6.1 and rebuilt and refined using WinCoot v0.9.8.7^79^. For mt-RNase Z^Gln^, the residues of pre-tRNA^Tyr^ were iteratively substituted to match the sequence of pre-tRNA^Gln^. The complete models of mt-RNase Z^Gln^ and mt-RNase Z^Tyr^ complexes were refined against the respective composite maps using ISOLDE^80^ and phenix.real-space refine^78,86^, followed by a single round of ADP refinement in PHENIX^78,86^ against the respective consensus maps. The MolProbity package within PHENIX suite was used for model validation^87^. The trajectory of the RNA in the productive substrate-engaged ELAC2 active site was based on the “contracted„ conformation of pre-mt-tRNA^Tyr^ **(Supplementary Figure 8D,E)**^80^. All structural analyses and image renderings for figure preparation were done using UCSF ChimeraX 1.6.1^81,88^ or PyMol 2.0 (The PyMOL Molecular Graphics System, Version 2.0 SchrÖdinger, LLC).

## DATA AVAILABILITY

The cryo-EM density reconstructions for mt-RNase Z^Gln^ and mt-RNase Z^Tyr^ were deposited with the Electron Microscopy Database (EMDB) under accession codes EMD-19455 and EMD-19457. The respective structure coordinates were deposited with the Protein Data Bank (PDB) under accession codes 8RR3 and 8RR4.

## ACKNOWLEDGEMENTS

We thank Christian Dienemann and Ulrich Steuerwald (MPI-NAT cryo-EM facility) for assistance with cryo-EM data acquisition, and Paula Prado for help with formatting the manuscript.

This work was funded by the Deutsche Forschungsgemeinschaft under Germany’s Excellence Strategy EXC 2067/1-390729940 (H.S.H), FOR2848 (P10, H.S.H.), SFB1190 (P23 to H.S.H.), SFB1565 (Project number 469281184, P13 to H.S.H., P12 to K.E.B.) and by the European Union (ERC Starting Grant MitoRNA, grant agreement no. 101116869, to H.S.H.). Views and opinions expressed are however those of the author(s) only and do not necessarily reflect those of the European Union or the European Research Council Executive Agency. Neither the European Union nor the granting authority can be held responsible for them.

## AUTHOR CONTRIBUTIONS

A.B. purified proteins, reconstituted mt-RNase Z^Tyr^ complex, collected and processed cryo-EM data, and built and analyzed both structural models, B.K. purified proteins and carried out biochemical assays, R.Y. purified proteins, reconstituted and prepared cryo-EM grids of mt-RNase Z^Gln^ and collected cryo-EM data, L.S. purified proteins and prepared cryo-EM grids of mt-RNase Z^Tyr^ together with A.B., K.D. performed insect cell culture and protein expression. H.S.H. conceived the study and supervised research. A.B., B.K., K.E.B. and H.S.H. wrote the manuscript.

## REFERENCES

1. Hoagland, M.B., Stephenson, M.L., Scott, J.F., Hecht, L.I., and Zamecnik, P.C. (1958). A soluble ribonucleic acid intermediate in protein synthesis. Journal of Biological Chemistry 231, 241–257.

2. Wolin, S.L., and Matera, A.G. (1999). The trials and travels of tRNA. Genes Dev 13, 1–10. 10.1101/GAD.13.1.1.

3. Deutscher, M.P. (1984). Processing of tRNA in prokaryotes and eukaryote. Crit Rev Biochem Mol Biol 17, 45–71. 10.3109/10409238409110269/ASSET//CMS/ASSET/A1BBFFFE-373D-410B-80EB-5035A5C70F9A/10409238409110269.FP.PNG.

4. Phizicky, E.M., and Hopper, A.K. (2015). tRNA processing, modification, and subcellular dynamics: past, present, and future. RNA 21, 483–485. 10.1261/RNA.049932.115.

5. Huxley, H.E., Brown, W.J., Elliott, G.F., Lowy, J., Millman, B.M.J., Rosenbaum, G., Holmes, K.C., Witz, J., Reynolds, G.T., Milch, J.R., et al. (1980). Order and intracellular location of the events involved in the maturation of a spliced tRNA. Nature 1980 284:5752 284, 143–148. 10.1038/284143a0.

6. Castaño, J.G., Tobian, J.A., and Zasloff, M. (1985). Purification and characterization of an endonuclease from Xenopus laevis ovaries which accurately processes the 3’ terminus of human pre-tRNA-Met(i) (3’ pre-tRNase). J Biol Chem 260, 9002–9008.

7. Li, Z., and Deutscher, M.P. (1996). Maturation pathways for E. coli tRNA precursors: A random multienzyme process in vivo. Cell 86, 503–512. 10.1016/S0092-8674(00)80123-3.

8. Stark, B.C., Kole, R., Bowman, E.J., and Altman, S. (1978). Ribonuclease P: an enzyme with an essential RNA component. Proceedings of the National Academy of Sciences 75, 3717–3721. 10.1073/PNAS.75.8.3717.

9. Schürer, H., Schiffer, S., Marchfelder, A., and Mörl, M. (2001). This is the end: Processing, editing and repair at the tRNA 3′-terminus. Biol Chem 382, 1147–1156. 10.1515/BC.2001.144/MACHINEREADABLECITATION/RIS.

10. Green, C.J., and Vold, B.S. (2014). tRNA, tRNA Processing, and Aminoacyl-tRNA Synthetases. Bacillus subtilis and Other Gram-Positive Bacteria, 683–698. 10.1128/9781555818388.CH47.

11. Suzuki, T. (2021). The expanding world of tRNA modifications and their disease relevance. Nature Reviews Molecular Cell Biology 2021 22:6 22, 375–392. 10.1038/s41580-021-00342-0.

12. Motorin, Y., and Helm, M. (2010). tRNA stabilization by modified nucleotides. Biochemistry 49, 4934–4944. 10.1021/BI100408Z.

13. Slade, A., Kattini, R., Campbell, C., and Holcik, M. (2020). Diseases Associated with Defects in tRNA CCA Addition. International Journal of Molecular Sciences 2020, Vol. 21, Page 3780 21, 3780. 10.3390/IJMS21113780.

14. Orellana, E.A., Siegal, E., and Gregory, R.I. (2022). tRNA dysregulation and disease. Nature Reviews Genetics 2022 23:11 23, 651–664. 10.1038/s41576-022-00501-9.

15. Haack, T.B., Kopajtich, R., Freisinger, P., Wieland, T., Rorbach, J., Nicholls, T.J., Baruffini, E., Walther, A., Danhauser, K., Zimmermann, F.A., et al. (2013). ELAC2 mutations cause a mitochondrial RNA processing defect associated with hypertrophic cardiomyopathy. Am J Hum Genet 93, 211–223. 10.1016/J.AJHG.2013.06.006.

16. Westhof, E., Thornlow, B., Chan, P.P., and Lowe, T.M. (2022). Eukaryotic tRNA sequences present conserved and amino acid-specific structural signatures. Nucleic Acids Res 50, 4100–4112. 10.1093/NAR/GKAC222.

17. Xue, H., Shen, W., Giegé, R., and Wong, J.T.F. (1993). Identity elements of tRNATrp: Identification and evolutionary conservation. Journal of Biological Chemistry 268, 9316–9322. 10.1016/s0021-9258(18)98352-3.

18. Hou, Y.M., Motegi, H., Lipman, R.S.A., Hamann, C.S., and Shiba, K. (1999). Conservation of a tRNA core for aminoacylation. Nucleic Acids Res 27, 4743–4750. 10.1093/NAR/27.24.4743.

19. Dudock, B.S., Katz, G., Taylor, E.K., and Holley, R.W. (1969). PRIMARY STRUCTURE OF WHEAT GERM PHENYLALANINE TRANSFER RNA*. Proceedings of the National Academy of Sciences 62, 941–945. 10.1073/PNAS.62.3.941.

20. Kim, S.H., Quigley, G.J., Suddath, F.L., Mcpherson, A., Sneden, D., Kim, J.J., Weinzierl, J., and Rich, A. (1973). Three-Dimensional Structure of Yeast Phenylalanine Transfer RNA: Folding of the Polynucleotide Chain. Science (1979) 179, 285–288. 10.1126/SCIENCE.179.4070.285.

21. Robertus, J.D., Ladner, J.E., Finch, J.T., Rhodes, D., Brown, R.S., Clark, B.F.C., and Klug, A. (1974). Structure of yeast phenylalanine tRNA at 3 Å resolution. Nature 1974 250:5467 250, 546–551. 10.1038/250546a0.

22. Giegé, R., Jühling, F., Pütz, J., Stadler, P., Sauter, C., and Florentz, C. (2012). Structure of transfer RNAs: similarity and variability. Wiley Interdiscip Rev RNA 3, 37–61. 10.1002/WRNA.103.

23. Zhang, J., and Ferré-D’amaré, A.R. (2016). The tRNA Elbow in Structure, Recognition and Evolution. Life 2016, Vol. 6, Page 3 6, 3. 10.3390/LIFE6010003.

24. Biela, A., Hammermeister, A., Kaczmarczyk, I., Walczak, M., Koziej, L., Lin, T.Y., and Glatt, S. (2023). The diverse structural modes of tRNA binding and recognition. Journal of Biological Chemistry 299, 104966. 10.1016/J.JBC.2023.104966.

25. Abe, T., Inokuchi, H., Yamada, Y., Muto, A., Iwasaki, Y., and Ikemura, T. (2014). TRNADB-CE: TRNA gene database well-timed in the era of big sequence data. Front Genet 5, 83271. 10.3389/FGENE.2014.00114/BIBTEX.

26. Parisien, M., Wang, X., and Pan, T. (2013). Diversity of human tRNA genes from the 1000-genomes project. RNA Biol 10, 1853–1867. 10.4161/RNA.27361.

27. Garone, C., Minczuk, M., D’Souza, A.R., and Minczuk, M. (2018). Mitochondrial transcription and translation: overview. Essays Biochem 62, 309–320. 10.1042/EBC20170102.

28. Suzuki, T., Nagao, A., and Suzuki, T. (2011). Human Mitochondrial tRNAs: Biogenesis, Function, Structural Aspects, and Diseases. Annu Rev Genet 45, 299–329. 10.1146/annurev-genet-110410-132531.

29. Helm, M., Brulé, H., Friede, D., Giegé, R., Pütz, D., and Florentz, C. (2000). Search for characteristic structural features of mammalian mitochondrial tRNAs. RNA 6, 1356–1379. 10.1017/S1355838200001047.

30. Kuhle, B., Chihade, J., and Schimmel, P. (2020). Relaxed sequence constraints favor mutational freedom in idiosyncratic metazoan mitochondrial tRNAs. Nature Communications 2020 11:1 11, 1–12. 10.1038/s41467-020-14725-y.

31. Rossmanith, W. (2011). Localization of Human RNase Z Isoforms: Dual Nuclear/Mitochondrial Targeting of the ELAC2 Gene Product by Alternative Translation Initiation. PLoS One 6, e19152. 10.1371/JOURNAL.PONE.0019152.

32. Holzmann, J., Frank, P., Löffler, E., Bennett, K.L., Gerner, C., and Rossmanith, W. (2008). RNase P without RNA: Identification and Functional Reconstitution of the Human Mitochondrial tRNA Processing Enzyme. Cell 135, 462–474. 10.1016/J.CELL.2008.09.013.

33. Kuhle, B., Hirschi, M., Doerfel, L.K., Lander, G.C., and Schimmel, P. (2023). Structural basis for a degenerate tRNA identity code and the evolution of bimodal specificity in human mitochondrial tRNA recognition. Nature Communications 2023 14:1 14, 1–13. 10.1038/s41467-023-40354-2.

34. Nagaike, T., Suzuki, T., Tomari, Y., Takemoto-Hori, C., Negayama, F., Watanabe, K., and Ueda, T. (2001). Identification and Characterization of Mammalian Mitochondrial tRNA nucleotidyltransferases. Journal of Biological Chemistry 276, 40041–40049. 10.1074/jbc.M106202200.

35. Kuhle, B., Hirschi, M., Doerfel, L.K., Lander, G.C., and Schimmel, P. (2022). Structural basis for shape-selective recognition and aminoacylation of a D-armless human mitochondrial tRNA. Nature Communications 2022 13:1 13, 1–12. 10.1038/s41467-022-32544-1.

36. Rossmanith, W., Tullo, A., Potuschak, T., Karwan, R., and Sbisà, E. (1995). Human mitochondrial tRNA processing. Journal of Biological Chemistry 270, 12885–12891. 10.1074/jbc.270.21.12885.

37. Jarrous, N., and Reiner, R. (2007). Human RNase P: a tRNA-processing enzyme and transcription factor. Nucleic Acids Res 35, 3519–3524. 10.1093/NAR/GKM071.

38. Bhatta, A., and Hillen, H.S. (2022). Structural and mechanistic basis of RNA processing by protein-only ribonuclease P enzymes. Trends Biochem Sci 47, 965–977. 10.1016/J.TIBS.2022.05.006.

39. Wu, J., Niu, S., Tan, M., Huang, C., Li, M., Song, Y., Wang, Q., Chen, J., Shi, S., Lan, P., et al. (2018). Cryo-EM Structure of the Human Ribonuclease P Holoenzyme. Cell 175, 1393–1404.e11. 10.1016/j.cell.2018.10.003.

40. Bhatta, A., Dienemann, C., Cramer, P., and Hillen, H.S. (2021). Structural basis of RNA processing by human mitochondrial RNase P. Nature Structural & Molecular Biology 2021 28:9 28, 713–723. 10.1038/s41594-021-00637-y.

41. Reinhard, L., Sridhara, S., and Hällberg, B.M. (2017). The MRPP1/MRPP2 complex is a tRNA-maturation platform in human mitochondria. Nucleic Acids Res 45, 12469–12480. 10.1093/nar/gkx902.

42. Brzezniak, L.K., Bijata, M., Szczesny, R.J., and Stepien, P.P. (2011). Involvement of human ELAC2 gene product in 3′ end processing of mitochondrial tRNAs. RNA Biol 8. 10.4161/rna.8.4.15393.

43. Schiffer, S., Rösch, S., and Marchfelder, A. (2002). Assigning a function to a conserved group of proteins: the tRNA 3′-processing enzymes. EMBO J 21, 2769–2777. 10.1093/EMBOJ/21.11.2769.

44. De La Sierra-Gallay, I.L., Pellegrini, O., and Condon, C. (2005). Structural basis for substrate binding, cleavage and allostery in the tRNA maturase RNase Z. Nature 2004 433:7026 433, 657–661. 10.1038/nature03284.

45. Ishii, R., Minagawa, A., Takaku, H., Takagi, M., Nashimoto, M., and Yokoyama, S. (2005). Crystal structure of the tRNA 3′ processing endoribonuclease tRNase Z from Thermotoga maritima. Journal of Biological Chemistry 280, 14138–14144. 10.1074/jbc.M500355200.

46. Schilling, O., Späth, B., Kostelecky, B., Marchfelder, A., Meyer-Klaucke, W., and Vogel, A. (2005). Exosite modules guide substrate recognition in the ZiPD/ElaC protein family. Journal of Biological Chemistry 280, 17857–17862. 10.1074/jbc.M500591200.

47. Li De La Sierra-Gallay, I., Mathy, N., Pellegrini, O., and Condon, C. (2006). Structure of the ubiquitous 3′ processing enzyme RNase Z bound to transfer RNA. Nature Structural & Molecular Biology 2006 13:4 13, 376–377. 10.1038/nsmb1066.

48. Vogel, A., Schilling, O., Späth, B., and Marchfelder, A. (2005). The tRNase Z family of proteins: physiological functions, substrate specificity and structural properties. Biol Chem 386, 1253–1264. 10.1515/BC.2005.142.

49. Wang, Z., Zheng, J., Zhang, X., Peng, J., Liu, J., and Huang, Y. (2012). Identification and Sequence Analysis of Metazoan tRNA 3′-End Processing Enzymes tRNase Zs. PLoS One 7, e44264. 10.1371/JOURNAL.PONE.0044264.

50. Takaku, H., Minagawa, A., Takagi, M., and Nashimoto, M. (2003). A candidate prostate cancer susceptibility gene encodes tRNA 3′ processing endoribonuclease. Nucleic Acids Res 31, 2272–2278. 10.1093/NAR/GKG337.

51. Jühling, F., Mörl, M., Hartmann, R.K., Sprinzl, M., Stadler, P.F., and Pütz, J. (2009). tRNAdb 2009: compilation of tRNA sequences and tRNA genes. Nucleic Acids Res 37. 10.1093/NAR/GKN772.

52. Suzuki, T., Yashiro, Y., Kikuchi, I., Ishigami, Y., Saito, H., Matsuzawa, I., Okada, S., Mito, M., Iwasaki, S., Ma, D., et al. (2020). Complete chemical structures of human mitochondrial tRNAs. Nature Communications 2020 11:1 11, 1–15. 10.1038/s41467-020-18068-6.

53. Zareen, N., Yan, H., Hopkinson, A., and Levinger, L. (2005). Residues in the Conserved His Domain of Fruit Fly tRNase Z that Function in Catalysis are Not Involved in Substrate Recognition or Binding. J Mol Biol 350, 189–199. 10.1016/J.JMB.2005.04.073.

54. Varadi, M., Anyango, S., Deshpande, M., Nair, S., Natassia, C., Yordanova, G., Yuan, D., Stroe, O., Wood, G., Laydon, A., et al. (2022). AlphaFold Protein Structure Database: massively expanding the structural coverage of protein-sequence space with high-accuracy models. Nucleic Acids Res 50, D439–D444. 10.1093/NAR/GKAB1061.

55. Chan, C.W., Chetnani, B., and Mondragón, A. (2013). Structure and function of the T-loop structural motif in noncoding RNAs. Wiley Interdiscip Rev RNA 4, 507–522. 10.1002/WRNA.1175.

56. Hingerty, B., Brown, R.S., and Jack, A. (1978). Further refinement of the structure of yeast tRNAPhe. J Mol Biol 124, 523–534. 10.1016/0022-2836(78)90185-7.

57. Qin, X., Deng, X., Chen, L., and Xie, W. (2016). Crystal Structure of the Wild-Type Human GlyRS Bound with tRNAGly in a Productive Conformation. J Mol Biol 428, 3603–3614. 10.1016/J.JMB.2016.05.018.

58. Ma, M., De La Sierra-Gallay, I.L., Lazar, N., Pellegrini, O., Durand, D., Marchfelder, A., Condon, C., and Van Tilbeurgh, H. (2017). The crystal structure of Trz1, the long form RNase Z from yeast. Nucleic Acids Res 45, 6209–6216. 10.1093/NAR/GKX216.

59. Pellegrini, O., Li De La Sierra-Gallay, I., Piton, J., Gilet, L., and Condon, C. (2012). Activation of tRNA Maturation by Downstream Uracil Residues in B. subtilis. Structure 20, 1769–1777. 10.1016/J.STR.2012.08.002.

60. Liao, R.Z., Himo, F., Yu, J.G., and Liu, R.Z. (2009). Theoretical Study of the RNA Hydrolysis Mechanism of the Dinuclear Zinc Enzyme RNase Z. Eur J Inorg Chem 2009, 2967–2972. 10.1002/EJIC.200900202.

61. Nashimoto, M., Tamura, M., and Kaspar, R.L. (1999). Selection of cleavage site by mammalian tRNA 3′ processing endoribonuclease. J Mol Biol 287, 727–740. 10.1006/JMBI.1999.2639.

62. Rackham, O., Busch, J.D., Matic, S., Siira, S.J., Kuznetsova, I., Atanassov, I., Ermer, J.A., Shearwood, A.-M.J., Richman, T.R., Stewart, J.B., et al. (2016). Hierarchical RNA Processing Is Required for Mitochondrial Ribosome Assembly. Cell Rep 16, 1874–1890. 10.1016/j.celrep.2016.07.031.

63. Watanabe, K. (2010). Unique features of animal mitochondrial translation systems. The non-universal genetic code, unusual features of the translational apparatus and their relevance to human mitochondrial diseases. Proc Jpn Acad Ser B Phys Biol Sci 86, 11–39. 10.2183/PJAB.86.11.

64. Wakita, K. Watanabe, Y. ichi, Yokogawa, T., Kumazawa, Y., Nakamura, S., Ueda, T., Watanabe, K., and Nishikawa, K. (1994). Higher-order structure of bovine mitochondrial tRNA Phe lacking the ‘conserved’ GG and TΨCG sequences as inferred by enzymatic and chemical probing. Nucleic Acids Res 22, 347–353. 10.1093/NAR/22.3.347.

65. Messmer, M., Pütz, J., Suzuki, T., Suzuki, T., Sauter, C., Sissler, M., and Catherine, F. (2009). Tertiary network in mammalian mitochondrial tRNAAsp revealed by solution probing and phylogeny. Nucleic Acids Res 37, 6881–6895. 10.1093/NAR/GKP697.

66. Vilardo, E., Toth, U., Hazisllari, E., Hartmann, R.K., and Rossmanith, W. (2023). Cleavage kinetics of human mitochondrial RNase P and contribution of its non-nuclease subunits. Nucleic Acids Res 51, 10536–10550. 10.1093/NAR/GKAD713.

67. Meynier, V., Hardwick, S.W., Catala, M., Roske, J., Oerum, S., Chirgadze, D.Y., Barraud, P., Yu, W., Luisi, B.F., and Tisné, C. (2023). Structural basis for human mitochondrial tRNA maturation. bioRxiv, 2023.12.19.572246. 10.1101/2023.12.19.572246.

68. Shi, P.Y., Maizels, N., and Weiner, A.M. (1998). CCA addition by tRNA nucleotidyltransferase: polymerization without translocation? EMBO J 17, 3197–3206. 10.1093/EMBOJ/17.11.3197.

69. Beuning, P.J., and Musier-Forsyth, K. (1999). Transfer RNA Recognition by Aminoacyl-tRNA Synthetases. Biopoly 52, 1–28. 10.1002/(SICI)1097-0282(1999)52:1.

70. Lehmann, J., Jossinet, F., and Gautheret, D. (2013). A universal RNA structural motif docking the elbow of tRNA in the ribosome, RNAse P and T-box leaders. Nucleic Acids Res 41, 5494–5502. 10.1093/NAR/GKT219.

71. D’Souza, A.R., Van Haute, L., Powell, C.A., Mutti, C.D., Páleníková, P., Rebelo-Guiomar, P., Rorbach, J., and Minczuk, M. (2021). YbeY is required for ribosome small subunit assembly and tRNA processing in human mitochondria. Nucleic Acids Res 49, 5798–5812. 10.1093/NAR/GKAB404.

72. Qin, X., Deng, X., Chen, L., and Xie, W. (2016). Crystal Structure of the Wild-Type Human GlyRS Bound with tRNAGly in a Productive Conformation. J Mol Biol 428, 3603–3614. 10.1016/J.JMB.2016.05.018.

73. Berger, I., Fitzgerald, D.J., and Richmond, T.J. (2004). Baculovirus expression system for heterologous multiprotein complexes. Nature Biotechnology 2004 22:12 22, 1583–1587. 10.1038/nbt1036.

74. Müller, M., Weigand, J.E., Weichenrieder, O., and Suess, B. (2006). Thermodynamic characterization of an engineered tetracycline-binding riboswitch. Nucleic Acids Res 34, 2607–2617. 10.1093/nar/gkl347.

75. Zivanov, J., Nakane, T., Forsberg, B.O., Kimanius, D., Hagen, W.J.H., Lindahl, E., and Scheres, S.H.W. (2018). New tools for automated high-resolution cryo-EM structure determination in RELION-3. Elife 7. 10.7554/ELIFE.42166.

76. Punjani, A., Rubinstein, J.L., Fleet, D.J., and Brubaker, M.A. (2017). CryoSPARC: Algorithms for rapid unsupervised cryo-EM structure determination. Nat Methods 14, 290–296. 10.1038/nmeth.4169.

77. Tegunov, D., and Cramer, P. (2019). Real-time cryo-electron microscopy data preprocessing with Warp. Nat Methods 16, 1146–1152. 10.1038/s41592-019-0580-y.

78. Liebschner, D., Afonine, P. V., Baker, M.L., Bunkoczi, G., Chen, V.B., Croll, T.I., Hintze, B., Hung, L.W., Jain, S., McCoy, A.J., et al. (2019). Macromolecular structure determination using X-rays, neutrons and electrons: recent developments in Phenix. urn: issn:2059-7983 75, 861–877. 10.1107/S2059798319011471.

79. Emsley, P., Lohkamp, B., Scott, W.G., and Cowtan, K. (2010). Features and development of Coot. Acta Crystallographica Section D 66, 486–501.

80. Croll, T.I. (2018). ISOLDE: A physically realistic environment for model building into low-resolution electron-density maps. Acta Crystallogr D Struct Biol 74, 519–530. 10.1107/S2059798318002425/IC5101SUP2.MP4.

81. Goddard, T.D., Huang, C.C., Meng, E.C., Pettersen, E.F., Couch, G.S., Morris, J.H., and Ferrin, T.E. (2018). UCSF ChimeraX: Meeting modern challenges in visualization and analysis. Protein Sci 27, 14–25. 10.1002/PRO.3235.

82. Gradia, S.D., Ishida, J.P., Tsai, M.S., Jeans, C., Tainer, J.A., and Fuss, J.O. (2017). MacroBac: New Technologies for Robust and Efficient Large-Scale Production of Recombinant Multiprotein Complexes. Methods Enzymol 592, 1–26. 10.1016/BS.MIE.2017.03.008.

83. Vos, S.M., Pöllmann, D., Caizzi, L., Hofmann, K.B., Rombaut, P., Zimniak, T., Herzog, F., and Cramer, P. (2016). Architecture and RNA binding of the human negative elongation factor. Elife 5. 10.7554/ELIFE.14981.

84. Kastner, B., Fischer, N., Golas, M.M., Sander, B., Dube, P., Boehringer, D., Hartmuth, K., Deckert, J., Hauer, F., Wolf, E., et al. (2008). GraFix: sample preparation for single-particle electron cryomicroscopy. Nat Methods 5, 53–55. 10.1038/NMETH1139.

85. Mastronarde, D.N. (2003). SerialEM: A program for automated tilt series acquisition on Tecnai microscopes using prediction of specimen position. In Microscopy and Microanalysis (Cambridge University Press), pp. 1182–1183. 10.1017/s1431927603445911.

86. Afonine, P. V., Poon, B.K., Read, R.J., Sobolev, O. V., Terwilliger, T.C., Urzhumtsev, A., and Adams, P.D. (2018). Real-space refinement in PHENIX for cryo-EM and crystallography. urn:issn:2059-7983 74, 531–544. 10.1107/S2059798318006551.

87. Williams, C.J., Headd, J.J., Moriarty, N.W., Prisant, M.G., Videau, L.L., Deis, L.N., Verma, V., Keedy, D.A., Hintze, B.J., Chen, V.B., et al. (2018). MolProbity: More and better reference data for improved all-atom structure validation. Protein Science 27, 293–315. 10.1002/PRO.3330.

88. Pettersen, E.F., Goddard, T.D., Huang, C.C., Meng, E.C., Couch, G.S., Croll, T.I., Morris, J.H., and Ferrin, T.E. (2021). UCSF ChimeraX: Structure visualization for researchers, educators, and developers. Protein Sci 30, 70–82. 10.1002/PRO.3943.

