## Supplementary Material for "Molecular basis of human nuclear and mitochondrial tRNA 3’-processing"

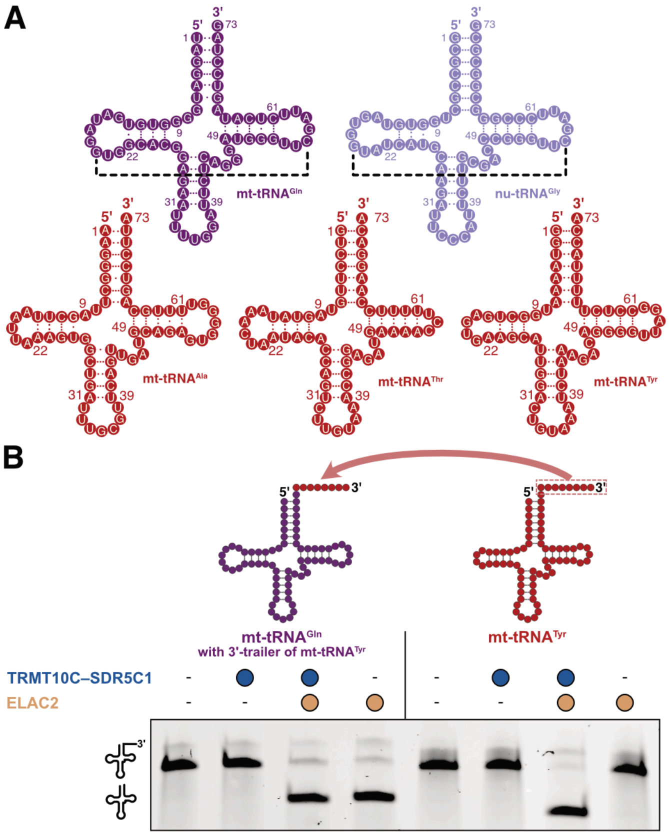


**Supplementary Figure 1 | Differential dependence of human tRNAs on TRMT10C–SDR5C1 for 3'-processing**
**A)** Schematic cloverleaf representation of all human tRNAs used in this study. Mt-tRNAs lacking the canonical elbow structures and requiring TRMT10C–SDR5C1 for 3'-processing: mt-tRNA$^{Ala}$, mt-tRNA$^{Thr}$ and mt-tRNA$^{Tyr}$, are shown in red; mt-tRNA$^{Gln}$ and nu-tRNA$^{Gly}$, which possess canonical elbow interactions and do not require TRMT10C–SDR5C1 for 3' processing, are shown in dark purple and light purple, respectively. G-C, A-U and G-U base pairs are indicated by three, two and one dots, respectively. Canonical G19-C56 base pairs are indicated by dashed black line segments.
**B)** *In vitro* cleavage assay showing that the differential dependence of mt-tRNA$^{Gln}$ and mt-tRNA$^{Tyr}$ on TRMT10C–SDR5C1, as observed in **Figure 1C**, does not relate to differences in the 3'-trailer sequences. The mt-tRNA$^{Gln}$ construct used in this assay contains the 3'-trailer sequence derived from mt-tRNA$^{Tyr}$.

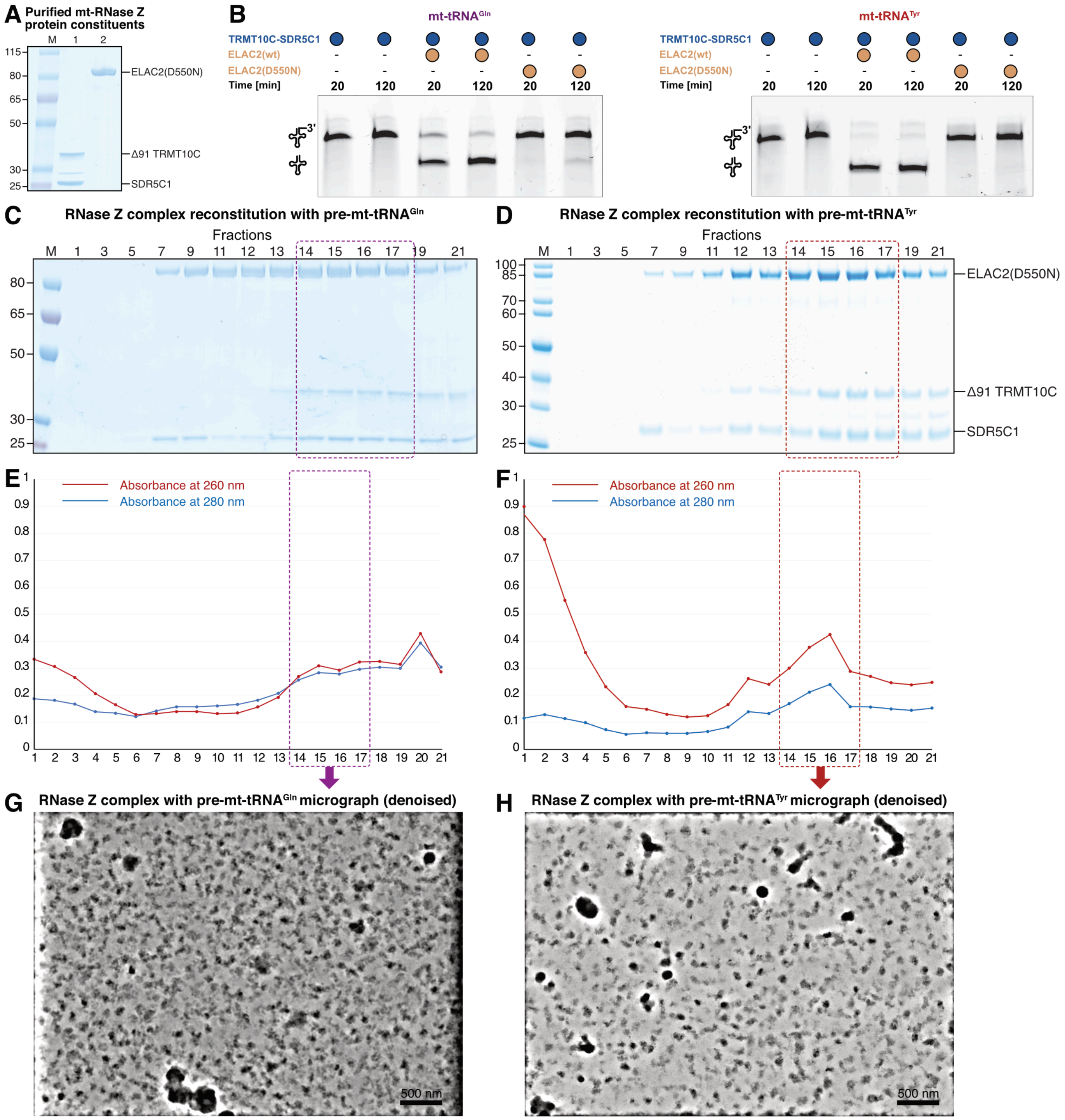


**Supplementary Figure 2 | *In vitro* reconstitution of mt-RNase Z complexes**

**A)** SDS-PAGE showing purified protein components of the mt-RNase Z complex.

**B)** *In vitro* cleavage assay showing the reduction in catalytic activity of the ELAC2(D550N) variant used for complex reconstitution.

**C)** SDS-PAGE analysis of the fractions of a representative sucrose density gradient ultracentrifugation (SDG) without glutaraldehylde crosslinker for mt-RNase $Z^{Gln}$. The sample used for cryo-EM was prepared in the presence of the crosslinker.

**D)** SDS-PAGE analysis of the fractions of a representative sucrose density gradient ultracentrifugation (SDG) without glutaraldehylde crosslinker for mt-RNase $Z^{Tyr}$. The sample used for cryo-EM was prepared in the presence of the crosslinker.

**E)** Line graph showing absorbance at 260 nm and 280 nm for SDG fractions for mt-RNase $Z^{Gln}$.

**F)** Line graph showing absorbance at 260 nm and 280 nm for SDG fractions for mt-RNase $Z^{Tyr}$.

**G)** A denoised representative cryo-EM micrograph of mt-RNase $Z^{Gln}$.

**H)** A denoised representative cryo-EM micrograph of mt-RNase $Z^{Gln}$.

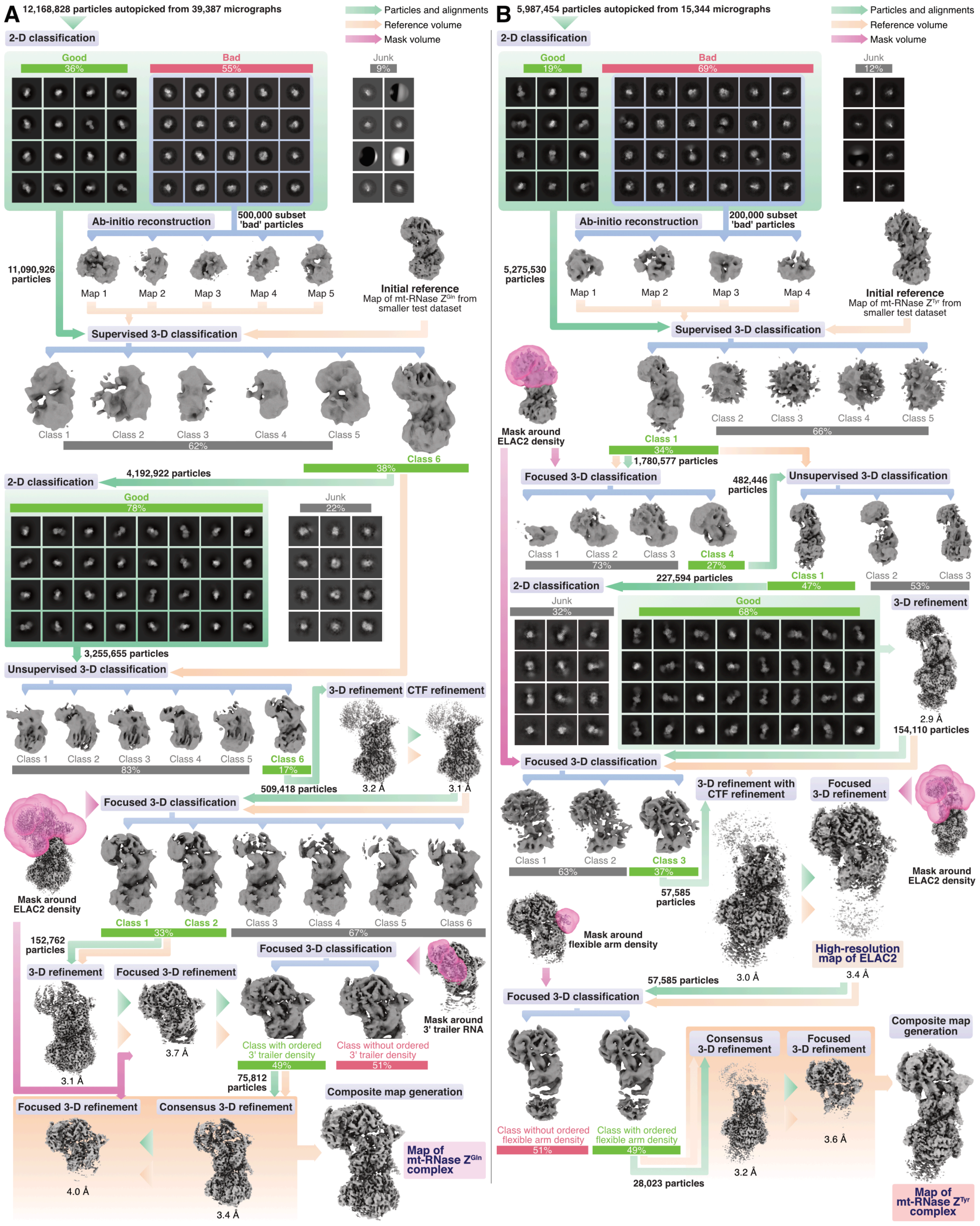


**Supplementary Figure 3 I Cryo-EM processing**

**A)** Cryo-EM processing workflow for mt-RNase $Z^{Gln}$ complex.

**B)** Cryo-EM processing workflow for mt-RNase $Z^{Tyr}$ complex

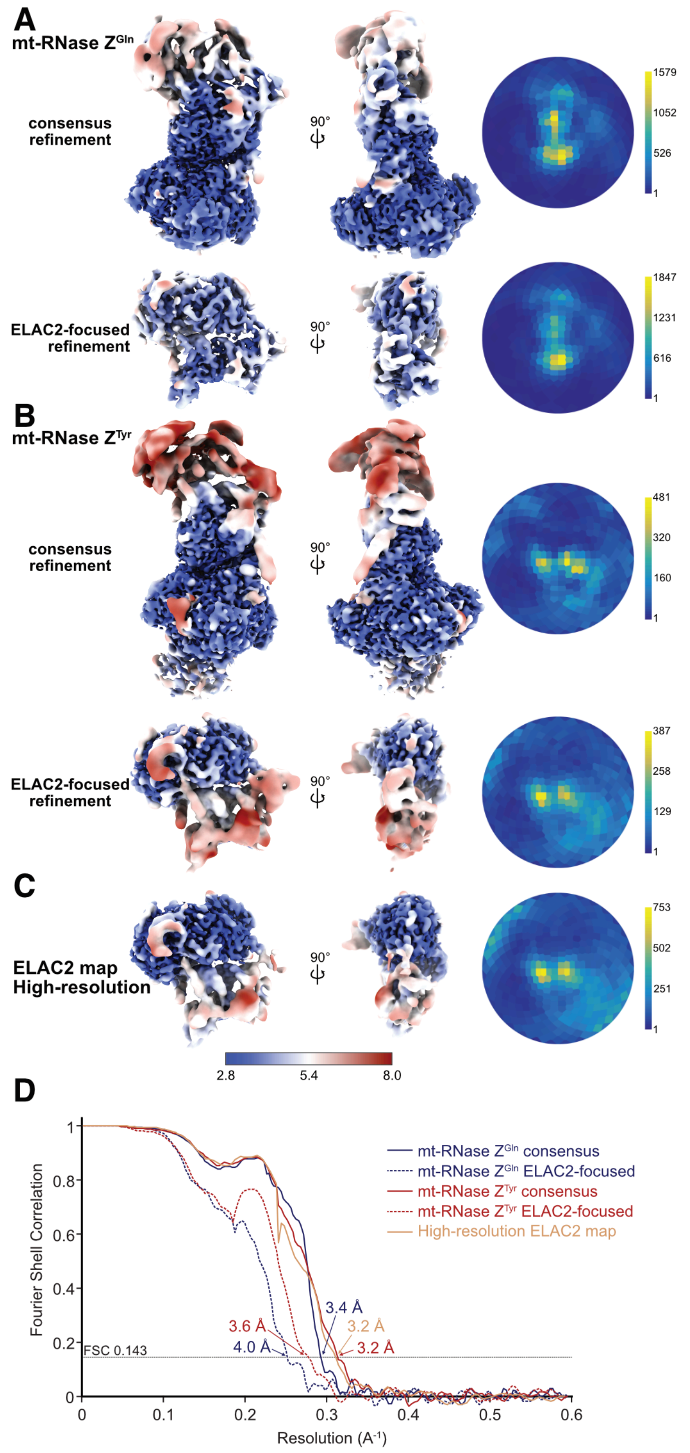


**Supplementary Figure 4 I Cryo-EM post-processing analysis**

**A)** Local resolution maps (left) and angular distribution plots (right) for consensus refinement map and ELAC2-focused map of mt-RNase $Z^{Gln}$.

**B)** Local resolution maps (left) and angular distribution plots (right) for consensus refinement map and ELAC2-focused map of mt-RNase $Z^{Tyr}$.

**C)** Local resolution map (left) and angular distribution plot (right) for high-resolution ELAC2-focused map from the mt-RNase $Z^{Tyr}$ dataset.

**D)** Fourier Shell Correlation (FSC) plots for maps in **A-D**. FSC threshold of 0.143 was used for reporting resolutions.

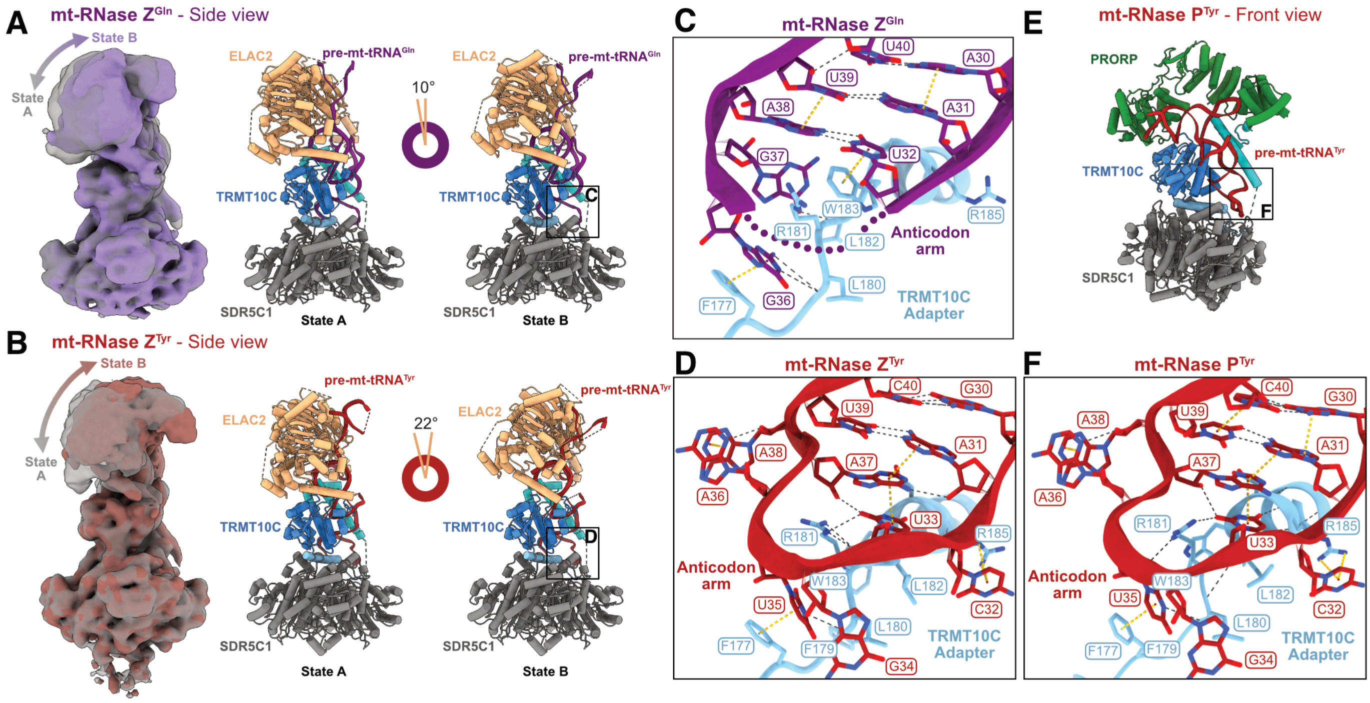


**Supplementary Figure 5 I Structural comparison of mt-RNase Z$^{Gln}$, mt-RNase Z$^{Tyr}$ and mt-RNase P$^{Tyr}$ complexes**

**A)** Conformational variability of ELAC2 in the mt-RNase Z$^{Gln}$ complex. States A and B represent two ends of a continuum of variability.

**B)** Conformational variability of ELAC2 in the mt-RNase Z$^{Tyr}$ complex. ELAC2 exhibits higher conformational variability in mt-RNase Z$^{Tyr}$ compared to mt-RNase Z$^{Gln}$.

**C)** Interactions between TRMT10C and the mt-tRNA$^{Gln}$ anticodon stem-loop in mt-RNase Z$^{Gln}$. The A38-U32 base pair extends the anticodon stem by one base pair and leads to a distinct stem-loop topology compared to mt-tRNA$^{Tyr}$ (**D,F**).

**D)** Interactions between TRMT10C and mt-tRNA$^{Tyr}$ anticodon stem-loop mt-RNase Z$^{Tyr}$. The anticodon stem-loop topology is similar to that previously observed in mt-RNase P$^{Tyr}$ (**E,F**)[40].

**E)** Structural model of mt-RNase P$^{Tyr}$ (PDB: 7ONU)[40].

**F)** Interactions between TRMT10C and the mt-tRNA$^{Tyr}$ anticodon stem-loop in mt-RNase P$^{Tyr}$.

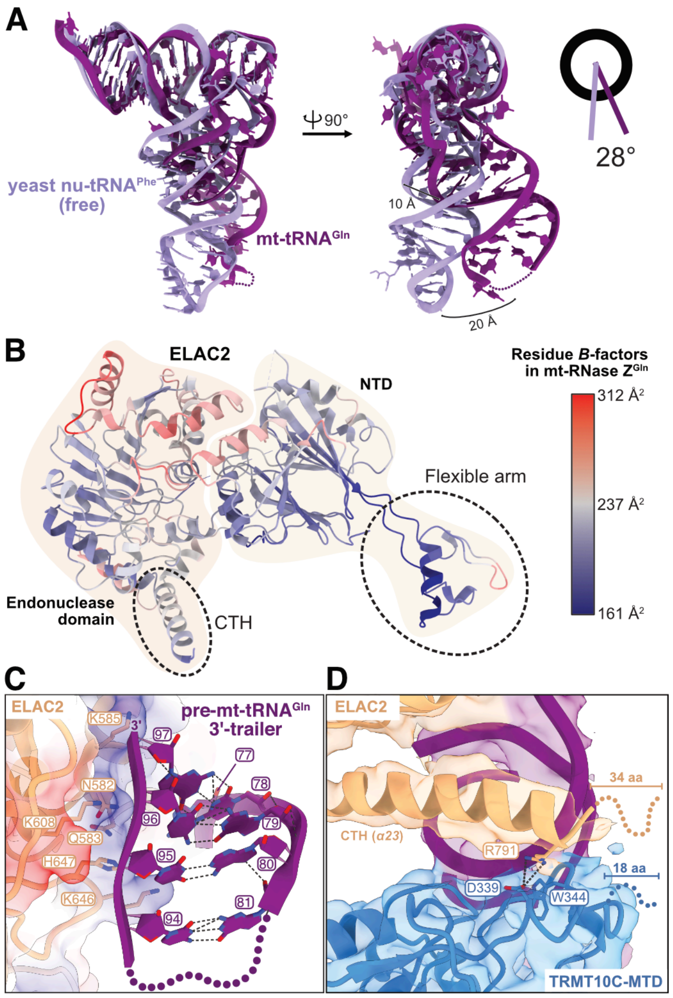


**Supplementary Figure 6 I Recognition of canonical pre-mt-tRNA by ELAC2**

**A)** Structural comparison of mt-RNase Z-bound human mt-tRNA$^{Gln}$ and free yeast nu-tRNA$^{Phe}$ (PDB: 4TNA)[56].

**B)** Cartoon representation of ELAC2 in the mt-RNase Z$^{Gln}$ complex colored by per-residue *B*-factors. The flexible arm residues have the lowest *B*-factor values among ELAC2 residues.

**C)** Interactions of the helical segment of the 3'-trailer of pre-mt-tRNA$^{Gln}$ with a positively charged patch in the ELAC2 endonuclease domain.

**D)** Interactions between ELAC2-CTH and TRMT10C-MTD in the mt-RNase Z$^{Gln}$ complex. 'aa' is short for amino acids.

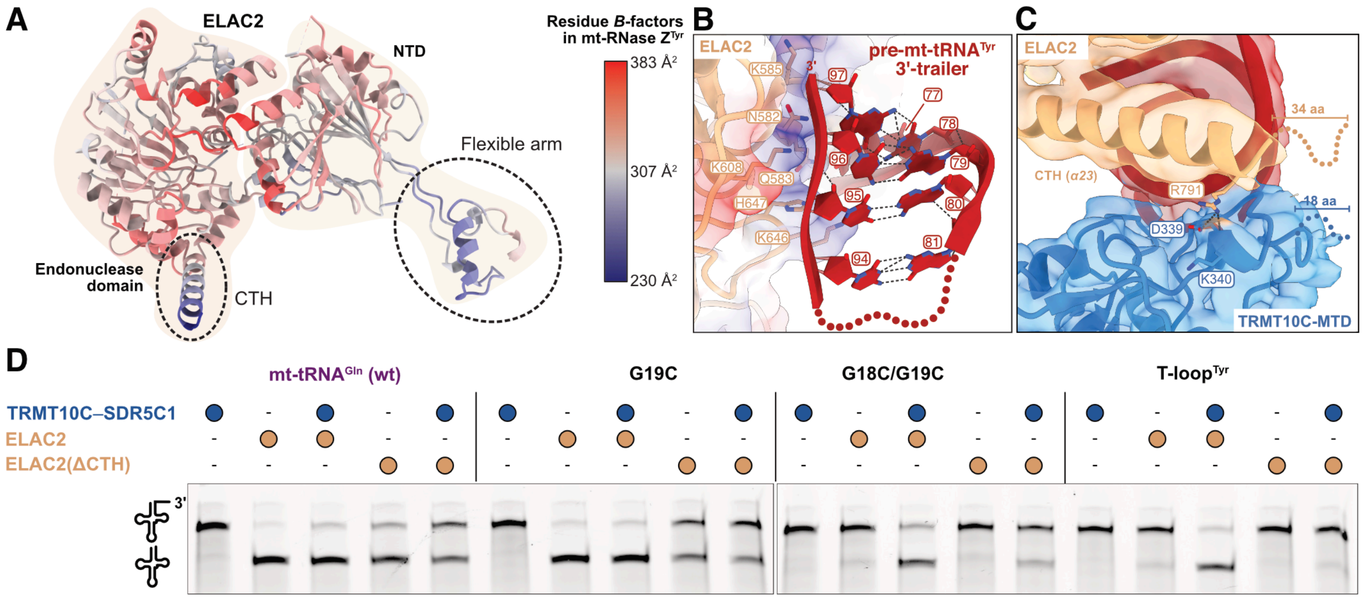


**Supplementary Figure 7 I Recognition of non-canonical pre-mt-tRNA by ELAC2**

**a)** Cartoon representation of ELAC2 in mt-RNase $Z^{Tyr}$ complex colored by per-residue *B*-factors. The flexible arm residues have lowest *B*-factor values among ELAC2 residues.

**b)** Interactions of the helical segment of the 3'-trailer of pre-mt-tRNA$^{Tyr}$ with a positively charged patch in the ELAC2 endonuclease domain.

**c)** Interactions between ELAC2-CTH and TRMT10C-MTD in mt-RNase $Z^{Tyr}$ complex.

**d)** *In vitro* cleavage assay demonstrating the role of ELAC2-CTH in 3'-processing of tRNAs. 3'-processing of mt-tRNA$^{Gln}$ variants with destabilized elbows is affected notably more than the wild-type mt-tRNA$^{Gln}$ with a stable canonical elbow.

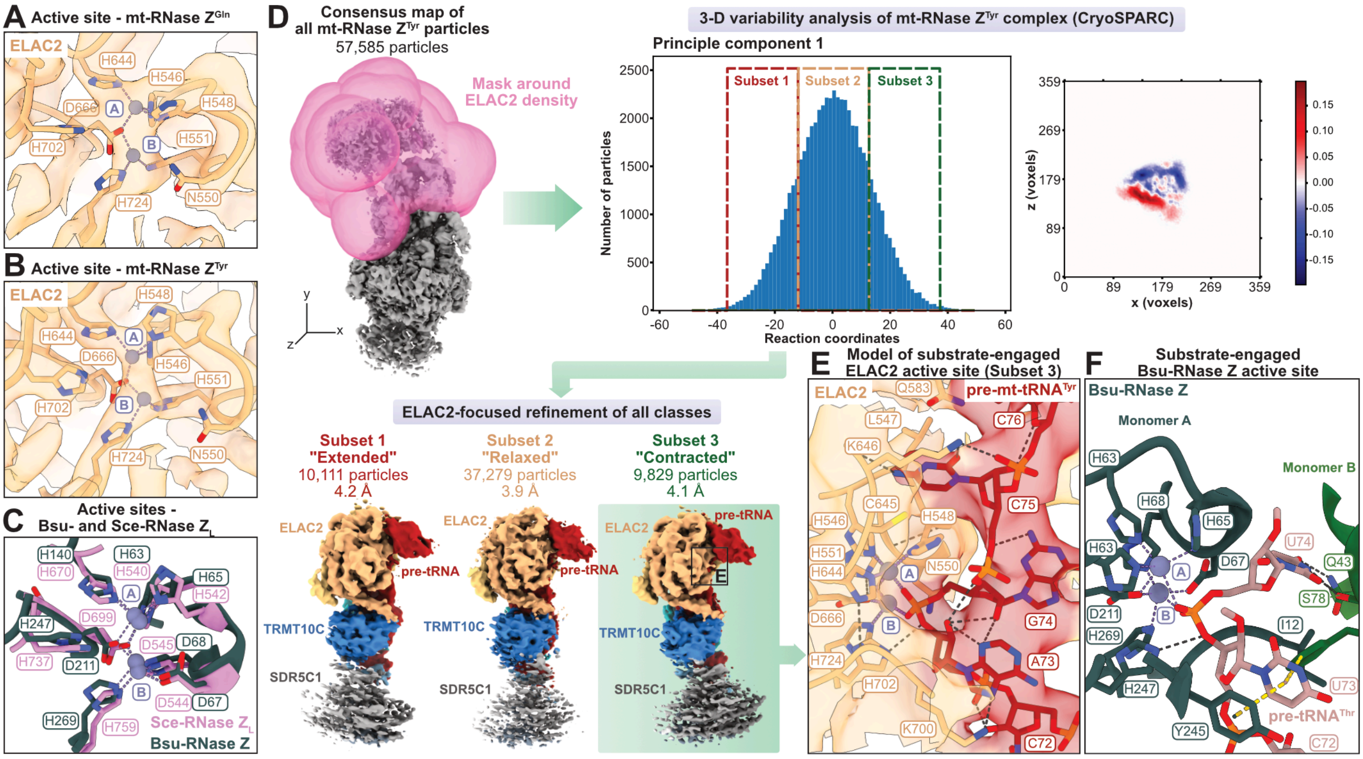


**Supplementary Figure 8 I Catalytic mechanism of ELAC2**

**A)** Active site of ELAC2 in the mt-RNase $Z^{Gln}$ complex. The cryo-EM density map is overlaid.

**B)** Active site of ELAC2 in mt-RNase $Z^{Tyr}$ complex. The cryo-EM density map is overlaid as in (**A**).

**C)** Active sites of *Bacillus subtilis (Bsu)*-RNase Z (PDB: 4GCW)[47] and *Saccharomyces cerevisiae (Sce)*-RNase $Z_L$ (PDB: 5MTZ)[58].

**D)** 3-D variability analysis of the mt-RNase $Z^{Tyr}$ complex. The “extended”, “relaxed” and “contracted” conformational states refer to the conformation of the pre-tRNA near the 3’-cleavage site. The structures shown here represent snapshots in a continuum of variability.

**E)** Model of the substrate-engaged ELAC2 active site based on the cryo-EM density map of the “contracted” state.

**F)** Structure of substrate-engaged *Bsu*-RNase Z active site (PDB: 4GCW)[59].

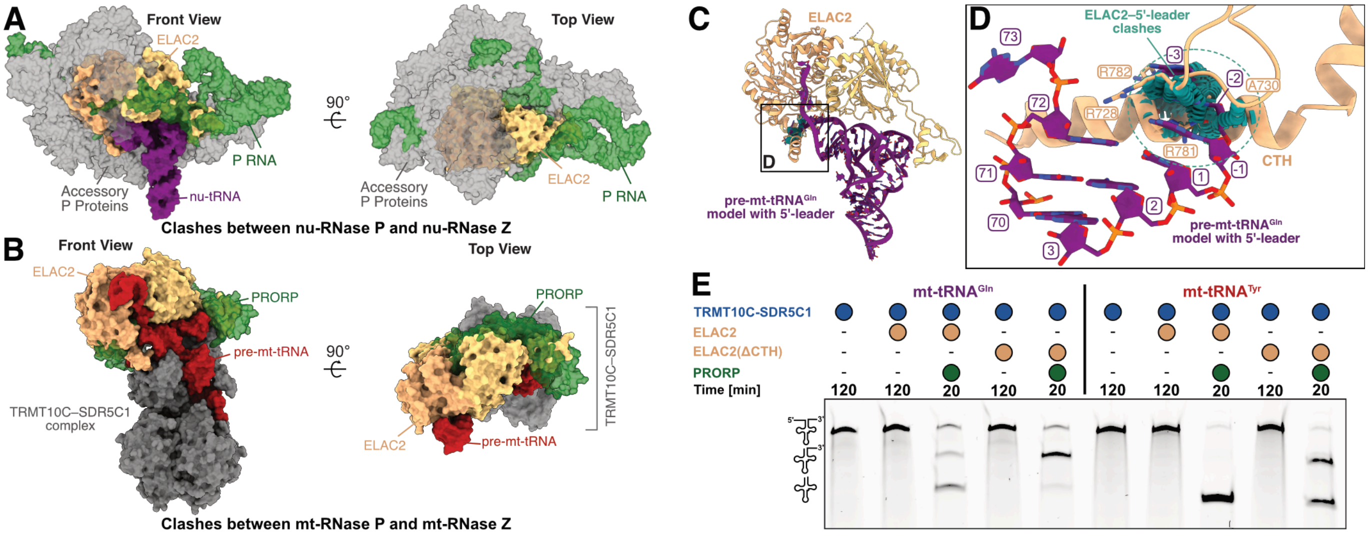


**Supplementary Figure 9 I Sequential order of tRNA maturation**

**A)** Overlay of nu-RNase P[39] and ELAC2 in complex with tRNA. Nu-RNase P is shown as transparent surface, with the P RNA shown in green and accessory proteins shown in grey. Binding of the two enzymes is mutually exclusive.

**B)** Overlay of the mt-RNase P[40] and mt-RNase Z complexes with tRNA. The PRORP subunit of mt-RNase P is shown as a transparent green surface. Binding of PRORP and ELAC2 to the TRMT10C–SDR5C1–pre-tRNA complex is mutually exclusive.

**C)** Structural model of ELAC2 with pre-mt-tRNA$^{Gln}$ showing clashes between ELAC2 and the 5'-leader. Region shown in detail in **D** is indicated.

**D)** Clashes between ELAC2 and modelled 5'-leader of pre-mt-tRNA$^{Gln}$. The view shown is rotated from **C** for clarity. Atom-to-atom clashes are shown in teal.

**E)** *In vitro* cleavage assay showing that the ELAC2-CTH is not the sole determinant of 5'–3' tRNA processing order.

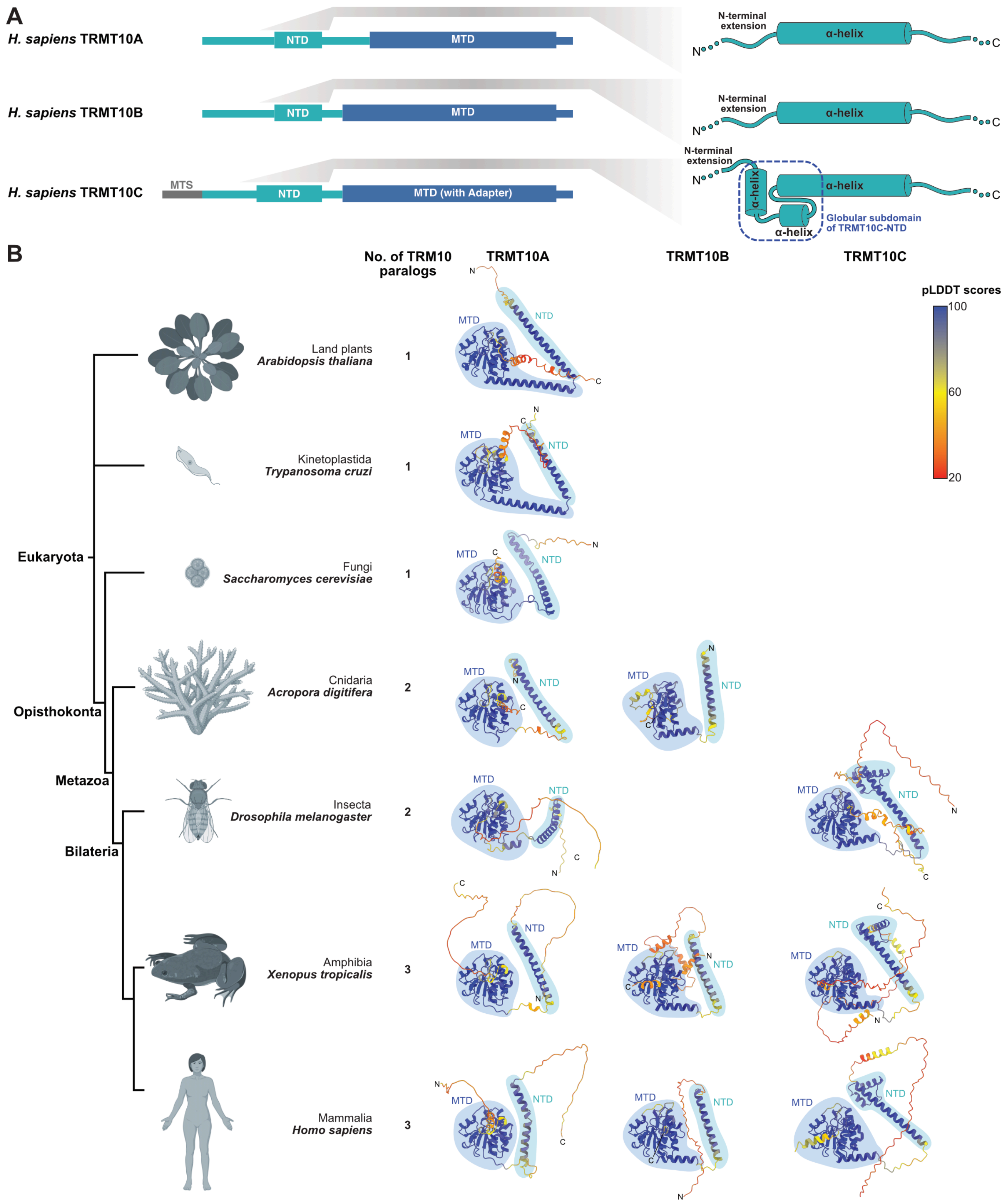


**Supplementary Figure 10 I TRM10 homologs across the eukaryotic domain of life**
**A)** Domain architectures of human TRM10 homologs. The NTD of TRMT10A and TRMT10B comprises a single long α-helix, while the TRMT10C-NTD is comprised of three α-helices that fold into a globular subdomain.
**B)** Comparison of Alphafold-predicted structures of TRM10 homologs across different eukaryotic taxa. The structures are colored by per-residue predicted local distance difference test (pLDDT) scores. The N-terminal domain (NTD) and methyltransferase domain (MTD) are marked and colored in cyan and blue, respectively. TRMT10C, with the globular subdomain in its NTD, is present only in bilaterian animals, coinciding with the structural erosion of their mitochondrial tRNAs[30]. Created with BioRender.com.
